# tRNA-derived second messengers confer bacterial immunity

**DOI:** 10.64898/2026.10.06.756102

**Authors:** Luis E. Valentin-Alvarado, Peter H. Yoon, Santiago C. Lopez, Yekaterina Shulgina, Conner Langeberg, Emily Armbruster, Muhammad Moez Ur-Rehman, Kenneth Loi, Cyntia Taveneau, Isabel Esain-Garcia, Mert Koka, Zeyuan Terry Zhang, Dmitry Kashin, Frank Schwede, Gaetan Burgio, Mitchell R. O’Connell, Erin E. Doherty, Jamie H. D. Cate, Gavin J. Knott, Jennifer A. Doudna

## Abstract

Bacterial small-molecule messengers that function in diverse immune signaling pathways have been thought to derive strictly from biosynthesis using nucleotide and cofactor substrates. Here we report a nuclease-driven immune signaling pathway in which tRNA cleavage generates a specific small molecule that triggers cell death in response to RNA-guided transcript recognition in prokaryotes. We identify a class of type VI-B CRISPR systems in which the RNA-guided nuclease Cas13 and its accessory transmembrane protein, 2TMβ, are co-encoded in an operon and are jointly required for bacteriophage defense. Upon target RNA recognition, Cas13 activates 2TMβ by producing mono- and dinucleotide messengers bearing 2′,3′-cyclic uridine phosphate ends by tRNA cleavage. Activated 2TMβ destroys infected cells by forming a multimeric complex that disrupts cell membranes to mediate cell death. These findings demonstrate RNA-programmed tRNA cleavage for second-messenger biogenesis, revealing a new logic for prokaryotic immune signaling.

## Introduction

Immune systems across all domains of life convert pathogen detection into diffusible chemical signals that activate cellular responses to infection. In bacteria, defense systems including CBASS, Thoeris, Pycsar, and type III CRISPR systems share a common logic in which infection triggers enzymatic production of a second messenger that binds to and activates downstream effectors^1–5^. Recent studies in prokaryotes have expanded the known types of signaling molecules^6–8^ and revealed that the signaling logic can be inverted to communicate safety rather than danger^9,10^. Despite this diversity, all known prokaryotic immune signaling pathways rely on dedicated enzymes that generate small-molecule signals from free nucleotides and cofactors^11^.

Eukaryotes generate small signaling molecules by processing larger precursors, including lipids and nucleic acids. During viral infection, the ribonuclease RNase L cleaves cellular and viral RNA into small fragments that activate the cytosolic receptor RIG-I to amplify the antiviral response^12^. During bacterial infection, the ribonuclease angiogenin cleaves the anticodon loops of tRNAs to generate tRNA halves that act as ligands for the endosomal Toll-like receptor 7^13^. Plant Toll/interleukin-1 receptor/resistance (TIR) domain containing receptors likewise hydrolyze nucleic acids to generate 2′,3′-cyclic mononucleotides that promote immune cell death^14^. In these cases, a nuclease that liberates signals from macromolecules, rather than an enzyme acting on a precursor pool, produces the signal. Whether the same strategy is employed in prokaryotic immune defense has remained unknown.

Comparative genomic analysis hinted that some prokaryotic defense systems might use a nuclease-based signaling strategy. RNA-guided RNA-targeting CRISPR-Cas13 systems can encode accessory genes that enhance immunity in a manner dependent on Cas13 nuclease activity, although how Cas13 activation is communicated to the accessory has been unresolved^15^. In this study, we identified Cas13 nucleases associated with the small trans-membrane protein 2TMβ, whose membrane-disrupting activity is regulated by binding to cyclic dinucleotides (CDNs) produced by nucleotidyltransferases in CBASS and Panoptes bacterial immune systems^9,10^. 2TMβ’s unexpected inclusion in some Cas13 operons implied an alternative mechanism of activation that might instead rely on nucleolytic activity.

Here we used a combination of genetic, biochemical and cell-based experiments to explore the relationship between Cas13 and 2TMβ, revealing a two-component nuclease-driven signaling pathway essential for viral defense. Both *in vivo* and *in vitro* experiments demonstrated that in a *2TMβ-cas13* system, RNA-guided transcript binding activates Cas13 as a tRNA-specific nuclease that generates cyclic-phosphate-containing products, including a guanylyl-(3′-5′)-uridine-2′,3′-cyclic monophosphate (GpU(2′,3′-cp)) linear dinucleotide and uridine-2′,3′-cyclic monophosphate (2′,3′-cUMP) mononucleotide. 2′,3′-cUMP and GpU(2′,3′-cp) bind to 2TMβ to promote its oligomerization into a membrane-perturbing structure that causes cell death. These results establish a new axis in bacterial immune signaling in which a nuclease-generated, tRNA-derived second messenger activates a dedicated effector to mount a defense response.

## Results

### A two-component type VI defense module

During investigation of diverse type VI CRISPR immune systems, which employ RNA-guided transcript recognition to initiate a response to viral infection, we noted some clades with colocalized genes encoding Cas13b and 2TMβ (**Fig. 1a, b; Extended Data Fig. 1a**). Two such representative loci, from *Chitinophaga sp*. and *Chryseobacterium sp*., encode deep-branching Cas13b effectors (Cas13b6 and Cas13b7) that define lineages we designated type VI-B-6 and VI-B-7; we refer to these two-gene systems as *2TMβ-cas13b6* and *2TMβ-cas13b7* (**Extended Data Fig. 1b**). Further genomic analysis revealed examples of 2TMβs encoded next to TIR domains or phosphodiesterases, raising the possibility that diverse enzymatic processes might generate ligands that regulate 2TMβ activity (**Extended Data Fig. 1c, d)**.

**Figure 1.**
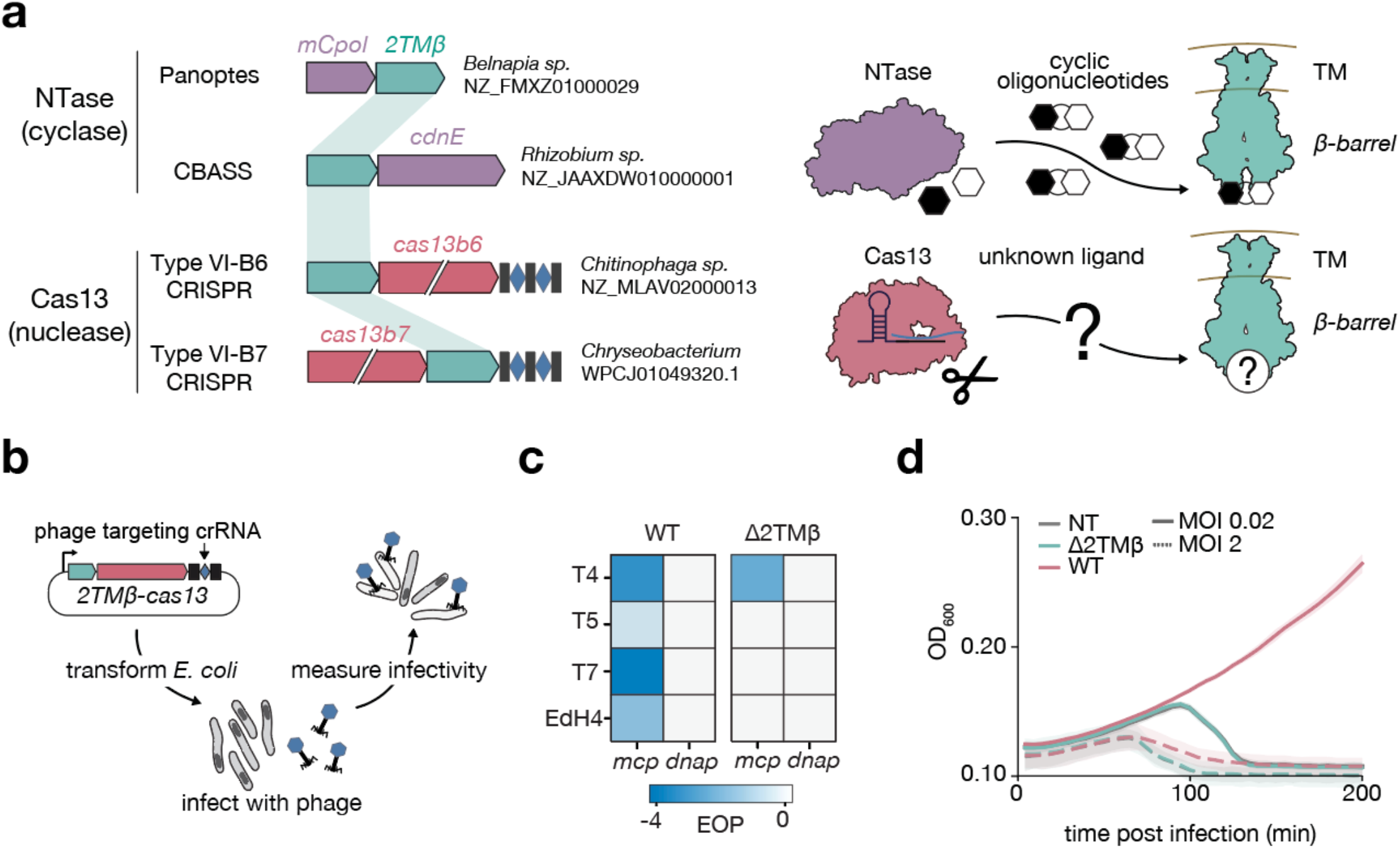
2TMβ-*cas13* is a two-component anti-phage module. **a**, Locus diagrams of 2TMβ-associated defense systems, spanning cyclase-based (Panoptes, CBASS) and nuclease-based (type VI-B6, VI-B7 CRISPR) systems (left). Models of the signaling pathways in the cyclase-based and Cas13-based systems (right). NTase, nucleotidyltransferase. **b**, Schematic of phage-defense assay in *E. coli*. **c**, Efficiency of plating (EOP, log10) for *dnap*- or *mcp*-targeting guides in 2TMβ-Cas13 (WT) or Δ2TMβ (Cas13-only) constructs. *dnap*, DNA polymerase gene; *mcp*, major capsid protein gene. **d**, Growth of non-target guide (NT), Δ2TMβ, and WT strains after T7 infection at multiplicity of infection (MOI) 0.02 and 2 (n = 3 biological replicates).

We tested whether *2TMβ-cas13* gene associations reflect a functional phage defense pathway by expressing the *2TMβ-cas13b6* and *2TMβ-cas13b7* systems bearing phage-targeting CRISPR RNAs (crRNAs) in *E. coli* and assessing their ability to provide immunity against a panel of lytic phages (**Fig. 1b, c**). Expression of the full *2TMβ-cas13b6* locus reduced the efficiency of plating by up to 4 orders of magnitude, and disruption of *2TMβ* abolished protection; the same pattern was also observed in the *2TMβ-cas13b7* system (**Fig. 1c; Extended Data Fig. 2a, b**). Quantification across multiple phages for *2TMβ-cas13b6* showed that robust defense required the complete system, with strong effects observed when targeting the major capsid protein (*mcp*) (**Fig. 1c**). Robust phage defense was observed in liquid infection assays: at low multiplicity of infection (MOI), cultures expressing the intact *2TMβ-cas13b6* locus were unaffected by phage infection, whereas cultures expressing a locus lacking *2TMβ* or expressing non-targeting guides collapsed (**Fig. 1d; Extended Data Fig. 3a-h**). Cas13b6 alone retained partial protection against T4 at low MOI (**Fig. 1c; Extended Data Fig. 3a-h**). At high MOI, all conditions collapsed, implying that *2TMβ-cas13b6* provides population-level immunity rather than directly interfering with the phage life cycle (**Fig. 1d; Extended Data Fig. 3d, h**). Together, these results establish *2TMβ-cas13b* as a two-component defense module.

### Cas13 cleavage generates cyclic-phosphate tRNA second messengers

We next asked whether Cas13b6 activity could trigger 2TMβ without phage infection. We reprogrammed the Cas13b6 guide RNA to target a plasmid-borne GFP transcript, leading to loss of cell viability upon GFP transcript binding if 2TMβ is a *bona fide* cell-death effector (**Fig. 2a**). In the presence of a targeting transcript, expression of the full *2TMβ-cas13b6* locus dramatically reduced colony formation, while expression of either gene alone did not affect cell viability (**Fig. 2a; Extended Data Fig. 4a, b)**. We further tested whether Cas13b6 nuclease activity underlies this response by mutating its Higher Eukaryotes and Prokaryotes Nucleotide-binding (HEPN) domain RNase catalytic residues, and found that these substitutions abolished the phenotype (**Fig. 2a**). Notably, Cas13b6 clade members universally carry a conserved histidine, H104, adjacent to the HEPN1 Rx_4_H catalytic motif in the RNase active site **(Fig. 2c)**. Alanine substitution of H104 phenocopied the HEPN domain catalytic residue mutations, suggesting that additional features beyond the typical HEPN RNase residues may be involved in substrate recognition and cleavage.

**Figure 2.**
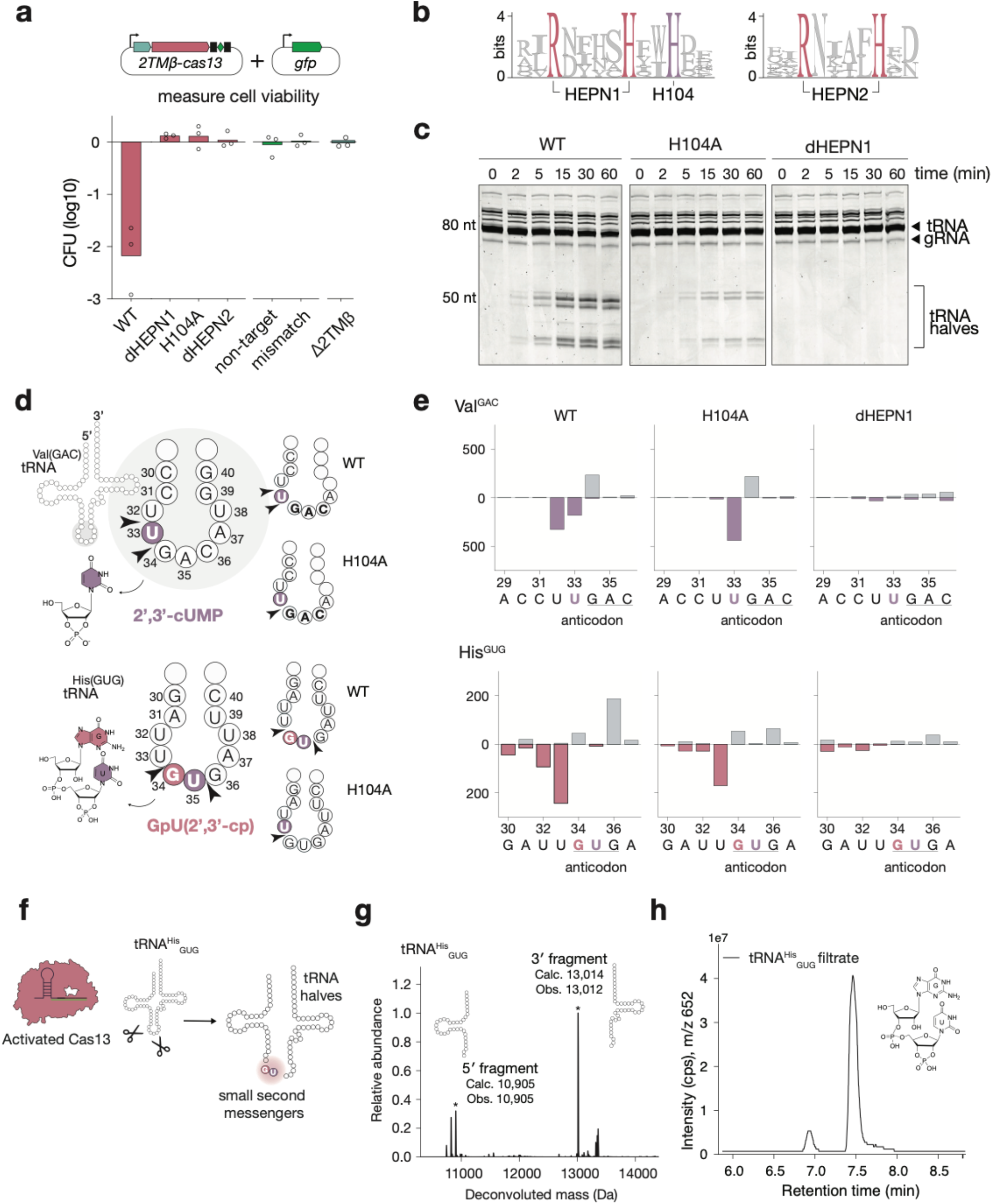
A conserved HEPN1-proximal histidine is required for paired anticodon-loop cleavage. **a**, Schematic of the phageless activation assay in *E. coli* (top). Viability of the indicated strains and guide controls, shown as colony-forming units (CFU, log_10_). Data points represent biological triplicates, bars indicate the mean (bottom). **b**, HEPN1 and HEPN2 region sequence logos in the Cas13b6 lineage. **c**, *In vitro* cleavage of purified *E. coli* tRNAs by WT Cas13b6, H104A, or dHEPN1, sampled at the indicated times in the presence of a complementary target RNA. **d**, Schematics of mapped anticodon-loop cleavage sites in tRNA^Val^_GAC_ and tRNA^His^_GUG_ for WT and H104A Cas13b6. Paired WT cleavage sites are consistent with excision of a single uridine from tRNA^Val^_GAC_ and a GU dinucleotide from tRNA^His^_GUG_; the corresponding candidate cyclic-phosphate products are shown. **e**, Small-RNA-seq profiles of Cas13-dependent 5’ and 3’ termini across the tRNA^Val^_GAC_ and tRNA^His^_GUG_ anticodon loops for WT, H104A and dHEPN1 Cas13b6. Grey and colored bars denote newly generated 5’ and 3’ termini, respectively. **f**, Schematic of Cas13b6 cleavage of *in vitro*-transcribed tRNA^His^_GUG_ and generation of complementary tRNA fragments and a short, excised product. **g**, Deconvoluted intact-RNA LC–MS spectrum of Cas13b6-treated, *in vitro*-transcribed tRNA^His^_GUG_. Asterisks mark the assigned 5’ and 3’ fragments. Calculated neutral masses (Calc.) and observed peak-apex masses (Obs.) are indicated to the nearest dalton. Calculations assume a 5’ triphosphate and terminal 2’,3’-cyclic phosphate for the 5’ fragment and hydroxyl termini for the 3’ fragment. Ion abundance is normalized to the maximum signal in the displayed mass range. **h**, Targeted LC–MS/MS analysis of the small-molecule filtrate from Cas13-treated IVT tRNA^His^_GUG_, detecting GpU(2’,3’-cp). dHEPN1: Cas13 R^96^NFYSH^101^ > A^96^NFYSA^101^; dHEPN2: Cas13 R^1106^NIALH^1111^ > A^1106^NIALA^1111^.

Since 2TMβ requires small nucleotide effector binding for activation in other prokaryotic immune systems ^9,10,16^, we wondered whether the native Cas13b6 RNase activity might itself be responsible for effector generation. We first examined Cas13b6 substrate preference using purified components. Target-activated Cas13b6 showed limited cleavage of a generic fluorogenic RNA reporter relative to LbuCas13a and RfxCas13d (**Extended Data Fig. 6a,b**). We therefore examined cleavage of cellular RNA substrates, beginning with purified *E. coli* tRNAs. Target-activated Cas13b6 generated discrete tRNA cleavage products (**Fig. 2c and Extended Data Fig. 7**). In reactions containing both tRNAs and a fluorescently labelled GFP target RNA, tRNA-derived products accumulated at early time points, whereas weak GFP target-RNA cleavage became apparent later (**Extended Data Fig. 6c**). Notably, Cas13b6 containing either HEPN1 or HEPN2 domain mutations abolished this activity, whereas H104A mutations impaired but did not abolish cleavage (**Fig. 2c**).

To identify the substrates and cleavage sites, we treated total E. coli RNA with target-activated Cas13b6 in vitro and mapped Cas13b6-dependent 5′ and 3′ RNA termini by small-RNA sequencing. Activation-associated termini mapped predominantly to tRNA anticodon loops, with cleavage after uridine (**Fig. 2d, e; Extended Data Figs. 8–9**). Separately, sequencing of RNA isolated from cells supported anticodon-region cleavage in vivo. Complementary MSR-seq of RNA from in vitro cleavage reactions recovered similar cleavage-associated patterns from tRNAs bearing modification-associated sequencing signatures (**Extended Data Figs. 10–13**). Analysis of the rRNA-rich fraction identified a single significantly enriched 3′ endpoint at uridine 2408 (U2408) in 23S rRNA (**Extended Data Fig. 14**). This pattern is reminiscent of activated LshCas13a and LbuCas13a, which cleave the uridine-rich anticodon loops in some tRNAs^17,18^.

sequence in tRNA^His^_GUG_ and tRNA^Asp^_GUG_, consistent with dinucleotide excision, whereas paired termini in tRNA^Val^_GAC_ flanked U33, consistent with single-uridine excision. H104A narrowed these paired-cleavage patterns and reduced the detected substrate range from 20 to 11 tRNAs, implicating H104 in short-product excision (**Fig. 2d, e; Extended Data Figs. 8,9,11,12)**. Activated Cas13b6 also cleaved isolated His and Val anticodon stem-loops, showing that the complete tRNA scaffold was not required for these substrates. Substitutions within the Val loop reduced the prominent cleavage-product signal, supporting a contribution of local sequence to cleavage **(Extended Data Fig. 15)**. Sequence-specific Northern blotting independently confirmed activator-dependent cleavage of His and Val tRNAs *in vitro* and *in vivo*, with no fragment accumulation in catalytically inactive dHEPN1 controls (dHEPN1: R^96^NFYSH^101^>A^96^NFYSA^101^; **Extended Data Figs. 16c–f and 17b–e)**. Cleavage persisted in Δ2TMβ cells, demonstrating that tRNA cleavage is independent of the membrane effector. At 2h post induction, fragments accounted for approximately 79–83% of the detected His and ValV Northern signal in this background, accompanied by reduced intact-tRNA signals (**Extended Data Fig. 17f,g**).

To determine the identities and terminal chemistry of the cleavage products, we analyzed *in vitro* transcribed and Cas13-treated His and Asp tRNAs by mass spectrometry. Intact-RNA LC–MS identified complementary fragments consistent with cleavage on either side of GU, with masses supporting a 2′,3′-cyclic phosphate at the 3′ end of the 5′ tRNA fragment the end chemistry (**Fig. 2f, g; Extended Data Fig. 18a-c**), expected of HEPN-family RNases^19^. To independently test the terminal chemistry, we compared 3′-adapter ligation before and after treatment with T4 polynucleotide kinase (PNK) that would restore 3′-hydroxyl groups compatible with adapter ligation. PNK treatment promoted ligation of the Cas13-derived fragments, and these ligated products migrated similarly to the ligation products of synthetic tRNA halves bearing 2′,3′-cyclic-phosphate termini **(Extended Data Fig. 19)**. This PNK dependence supports blocked 3′ termini, consistent with the cyclic-phosphate assignments from intact-RNA LC–MS. Targeted LC– MS/MS of small-molecule filtrates from *in vitro* Cas13b6 cleavage reactions with either tRNA^Asp^_GUG_ or tRNA^His^_GUG_ directly detected GpU(2′,3′-cp) by comparison with a synthetic standard, establishing release of a linear GU dinucleotide bearing a terminal uridine 2′,3′-cyclic phosphate **(Fig. 2h; Extended Data Fig. 18a, d)**.

Together, these findings demonstrate that Cas13b6 preferentially cleaves tRNA anticodon loops in *trans* to generate tRNA-derived second messengers bearing a terminal uridine 2′,3′-cyclic-phosphate moiety, including GpU(2′,3′-cp) and the candidate mononucleotide 2′,3′-cUMP **(Fig. 2d, e)**. H104A narrows the paired-cleavage pattern required for short-product excision and abolishes transcript-triggered toxicity in the phageless assay, identifying GpU(2′,3′-cp) and 2′,3′-cUMP as candidate signaling molecules in this system.

### Terminal uridine 2′,3′-cyclic-phosphate bearing ligands activate 2TMβ

The preceding results raised the possibility that Cas13-generated ligands activate 2TMβ directly. CBASS Cap15 and Panoptes immune systems also include 2TMβ, whose β-barrel domain recognizes the cyclase-generated ligand in its binding pocket^9,10,16^. Ligand binding leads to 2TMβ multimerization through direct barrel-barrel contacts^9,10,16^. The *2TMβ-cas13* β-barrel shares similar pocket organization with the CBASS and Panoptes 2TMβ, including the conserved tyrosine residues proposed to coordinate the ligand **(Fig. 3a)**. Motivated by this conservation, we tested whether the 2TMβ binding pocket recognizes Cas13b6-derived products. We assayed which components of the 2TMβ are essential for function using phage-free GFP-targeting assays **(Fig. 3b)**. We found that both the transmembrane domain and the β-barrel regions of 2TMβ are required for transcript-activated triggering of the *2TMβ-cas13b6* system **(Fig. 3b)**. In addition, Y157 and Y197 are both involved in this response, as alanine substitutions at these positions reduced activity **(Fig. 3b)**. These results are consistent with a ligand binding the 2TMβ pocket to drive signaling.

**Figure 3.**
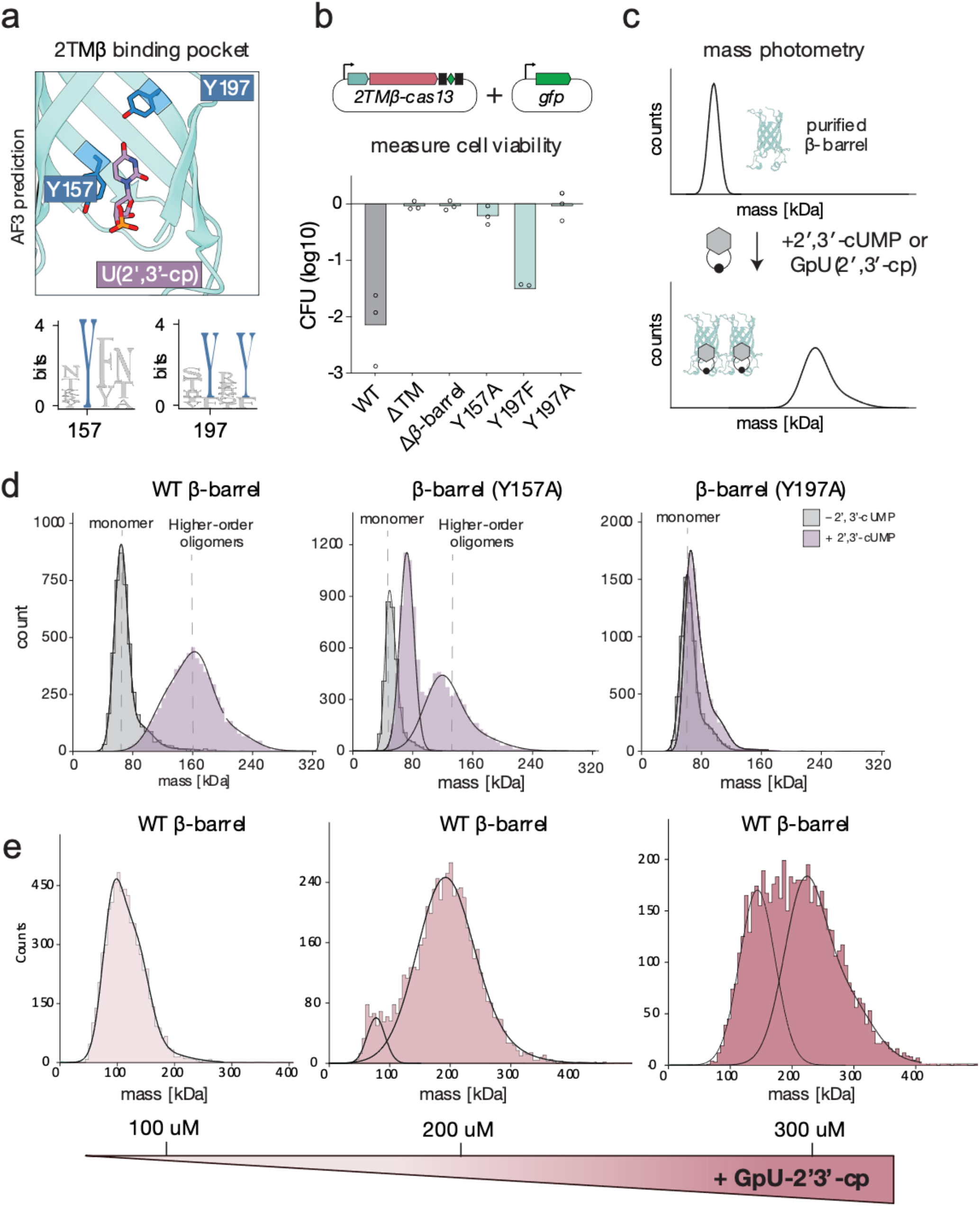
Uridine 2’,3’-cyclic-phosphate ligands promote 2TMβ assembly. **a**, Structure prediction of the 2TMβ β-barrel with 2’,3’-cUMP in the predicted ligand-binding pocket, highlighting Y157 and Y197 (top). Sequence logos show conservation of these positions across cas13-associated 2TMβ homologues (bottom). **b**, Phage-free GFP-targeting assay measuring colony formation in *E. coli* expressing Cas13b6 with wild-type 2TMβ or the indicated domain-deletion and pocket variants. Points represent three independent biological replicates; bars indicate the mean. **c**, Schematic of the ligand-response assay. Purified soluble 2TMβ β-barrel proteins were incubated with 2’,3’-cUMP or GpU(2’,3’-cp) and analyzed by mass photometry. **d**, Mass-photometry distributions of wild-type, Y157A and Y197A MBP–β-barrel proteins without (grey) or with (purple) 2’,3’-cUMP. The monomer reference and higher-order oligomer populations are indicated. **e**, Apparent mass distributions of MBP–β-barrel following incubation with increasing GpU(2’,3’-cp) concentrations. Concentrations shown refer to incubation mixtures containing 1 μM protein and 100, 200 or 300 μM ligand. Before measurement, 1 μl of each reaction was added to 19 μl of ligand-free buffer, yielding 50 nM protein and 5, 10 or 15 μM ligand, respectively. Each condition was tested in two independently assembled reactions (n = 2). In d,e, histograms show detected particle counts, with fitted distributions overlaid.

We next tested whether candidate ligands promote assembly of the purified 2TMβ barrel domain *in vitro*. To determine whether binding alters the assembly state of the protein, we incubated the purified β-barrel with 2′,3′-cUMP and a panel of ligands and analyzed these reactions by mass photometry **(Fig. 3c)**. Synthetic 2′,3′-cUMP shifted the predominantly monomeric WT protein towards higher-mass assemblies (**Fig. 3d**), with an independent titration revealing a concentration-dependent response (**Extended Data Fig. 20)**. GpU(2′,3′ cp) also promoted assembly, with higher-mass populations emerging as ligand concentration increased (**Fig. 3e**). These populations had higher apparent masses than those observed with 2′,3′-cUMP under the conditions tested, suggesting that ligand identity may influence the distribution of assembly sizes. Importantly, the pocket substitutions also altered ligand-responsive assembly. Y157A retained a reduced response to 2′,3′-cUMP, whereas Y197A did not produce a comparable higher-mass shift, supporting a role for the conserved pocket in ligand recognition (**Fig. 3d)**.

To examine the chemical requirements for assembly, we compared 2′,3′-cUMP with related nucleotides. All tested 2′,3′-cyclic mononucleotides (2′,3′cAMP, 2′,3′-cGMP and 2′,3′-cCMP) and cyclic oligonucleotides did not produce comparable higher-mass distributions (**Extended Data Fig. 21a)**. Likewise, β-Barrel oligomerization was unresponsive to the CDNs made by CBASS, type III CRISPR-Cas, and Panoptes cyclases (**Extended Data Fig. 22)**^9,10,16^. β-Barrel oligomerization was end-chemistry specific, as closely related forms such order state induced by 2′,3′-cUMP or GpU(2′,3′-cp) (**Extended Data Fig. 22**). Complementary structural predictions yielded higher confidence and more consistent configurations for 2′,3′-cUMP than for the other tested cyclic mononucleotides, providing a structural hypothesis for this preference **(Extended Data Fig. 21b, c)**.

These results establish that both 2′,3′-cUMP and GpU(2′,3′-cp) are sufficient to drive oligomerization of the isolated, soluble β-barrel *in vitro*. Together, these results support a model in which tRNA-derived second messengers bearing a terminal uridine 2′,3′-cyclic-phosphate moiety couple Cas13-mediated tRNA cleavage to 2TMβ assembly.

### Membrane lesions execute 2TMβ immunity

Having established this communication between Cas13 and 2TMβ both *in vitro* and *in vivo*, we next asked how 2TMβ activation mediates immunity. We monitored the kinetics of cell growth during phage-free GFP-targeting assays in liquid culture and found that expression of *2TMβ-cas13* causes rapid growth arrest **(Fig. 4a)**. Intracellular ATP was reduced in these arrested cells, likely due to ablation of the proton motive force caused by 2TMβ-induced inner membrane disruption **(Fig. 4b)**. Consistent with this, we found a substantial increase in membrane depolarization in arrested cells (**Extended Data Fig. 23**).

**Figure 4.**
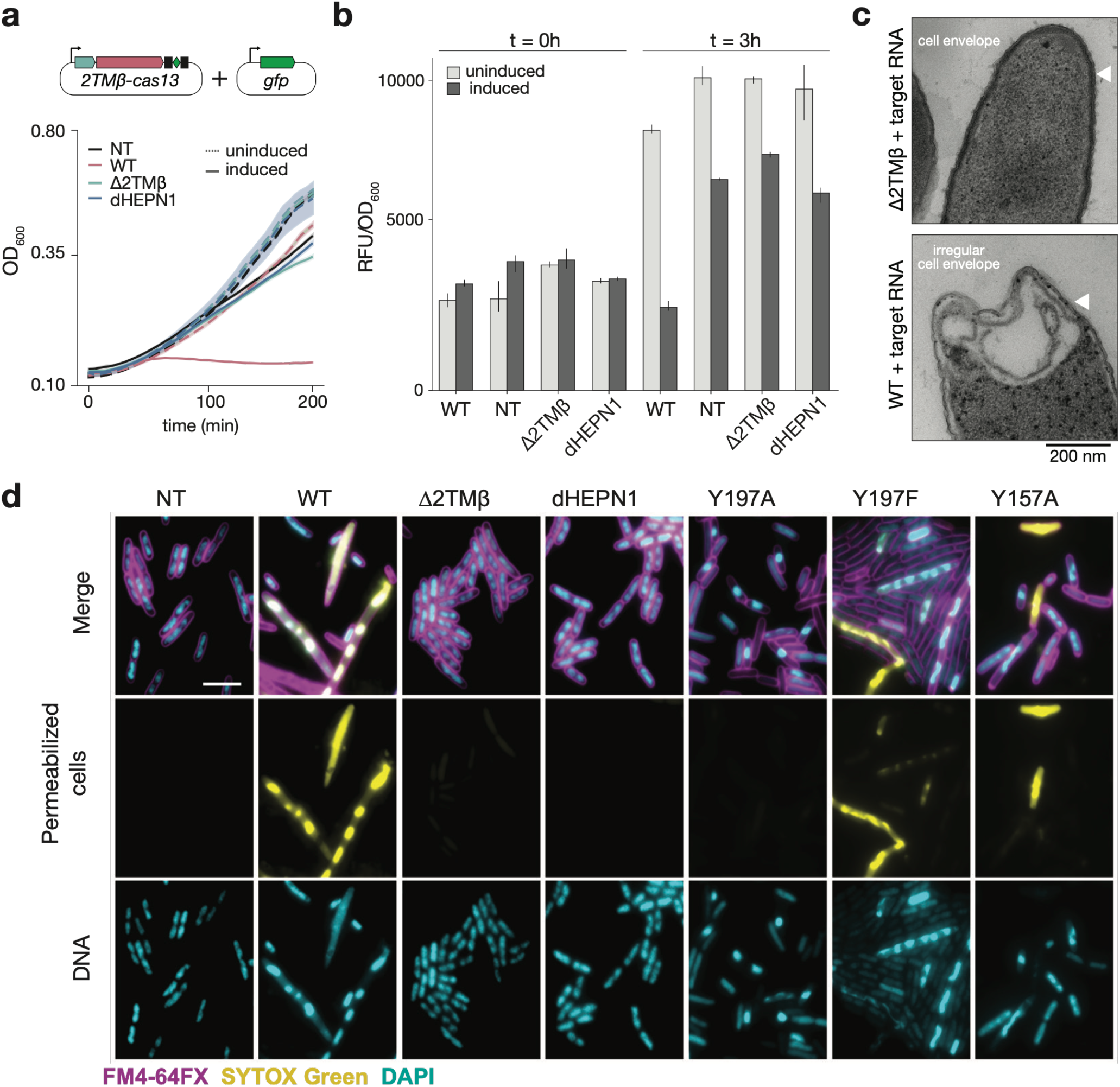
Activation of *2TMβ–cas13* causes growth arrest and membrane lesions. **a**, Schematic of the phage-free GFP-targeting assay in *E. coli* carrying the *2TMβ–cas13* locus and a *gfp* target plasmid (top), and growth of the indicated strains and guide controls (bottom). Solid and dotted lines denote induced and uninduced cultures, respectively. **b**, ATP-associated signal normalized to OD_600_ at 0 and 3 h in induced and uninduced cultures. Bars show the mean and error bars indicate s.d. **c**, Representative transmission electron microscopy images of Δ2TMβ and WT cells under targeting conditions. Scale bars, 200 nm. White arrowheads highlight cell-envelope morphology. **d**, Representative fluorescence microscopy images during the phage-free GFP-targeting assay. Cells were labelled with the membrane stain FM4-64FX (magenta), the membrane-permeabilization marker SYTOX Green (yellow) and the DNA stain DAPI (cyan). Scale bar, 5 μm. WT, complete *2TMβ–cas13b6* system with a targeting guide; NT, non-targeting guide; Δ2TMβ, deletion of the membrane effector; dHEPN1, Cas13b6 carrying alanine substitutions of the catalytic arginine and histidine in the HEPN1 RxxxxH motif (R^96^NFYSH^101^ > A^96^NFYSA^101^)

To distinguish the consequences of tRNA cleavage from those requiring 2TMβ, we compared polysome profiles. Despite detectable tRNA cleavage, Δ2TMβ cells retained prominent 70S and polysome peaks, showing that cleavage occurred without marked disruption of ribosome profiles at the sampled time point. These profiles were disrupted upon activation of the complete system, consistent with broader cellular damage associated with 2TMβ activity (**Extended Data Fig. 24**). Thus, tRNA cleavage occurred without comparable disruption of ribosome profiles in the absence of 2TMβ. In CBASS and

Panoptes, the 2TMβ effector is proposed to kill the cell by collapsing the membrane^9,10,16^. The ligand-responsive assembly of the isolated 2TMβ domain prompted us to examine membrane integrity in activated cells. Transmission electron microscopy revealed polar electron-lucent spaces and irregular envelope profiles in activated WT cells, contrasting with the more uniform appearance of the Δ2TMβ, non-targeting and dHEPN1 controls (**Fig. 4c; Extended Data Fig. 25**). Fluorescence microscopy revealed heterogeneous cell morphologies, including elongated cells, and pronounced uptake of the membrane-impermeable dye SYTOX Green in a subset of cells expressing the complete system, indicating membrane damage. Non-targeting, Δ2TMβ and HEPN-deficient controls showed little uptake (**Fig. 4d**).

To demonstrate that 2TMB ligand generation drives membrane perturbation, we assessed 2TMβ pocket substitutions *in vivo*. Consistent with their differences in oligomerization in response to 2′,3′-cUMP *in vitro*, these 2TMβ pocket mutants displayed a spectrum of growth arrest, cell morphology, and antiphage defense phenotypes (**Fig. 4d; Extended Data Fig. 5a, b**). Y157A retained detectable membrane-permeabilization activity and strong cell arrest phenotype, whereas Y197A was strongly impaired and displayed a reduced cell arrest phenotype, consistent with their different responses to 2′,3′-cUMP *in vitro*. The conservative Y197F substitution also retained SYTOX Green uptake (**Fig. 4d**). These results support a functional contribution of the ligand-responsive pocket to 2TMβ-dependent membrane damage. Together with the reduction in colony formation during phage-free activation (**Fig. 2a**), these observations support membrane damage and loss of viability as 2TMβ-dependent consequences of Cas13 activation.

## Discussion

Collectively, our results reveal a signaling pathway that connects RNA recognition to cell death. A guide RNA directs Cas13b6 to a target transcript, activating nucleolytic cleavage of a subset of host tRNA anticodon loops, releasing terminal U(2’,3’-cp) bearing ligands, including GpU(2′,3′-cp) and 2′,3′-cUMP. These ligands bind the β-barrel pocket of 2TMβ and drive its oligomerization to form membrane lesions, collapsing the cell’s energy supply and arresting its growth (**Fig. 5a**). The infected cell is killed, denying the phage a productive host, a logic that explains why protection holds at low MOI but is overwhelmed at high MOI (**Fig. 1d**). The *2TMβ-cas13b6* system exemplifies a nuclease-mediated immune response whose individual elements – tRNA as the source of the signal ^20^ and a small nucleotide released from a larger precursor as the messenger ^14^ – each have eukaryotic precedents. These similarities underscore the evolutionary parallels of immunity across the domains of life.

**Figure 5.**
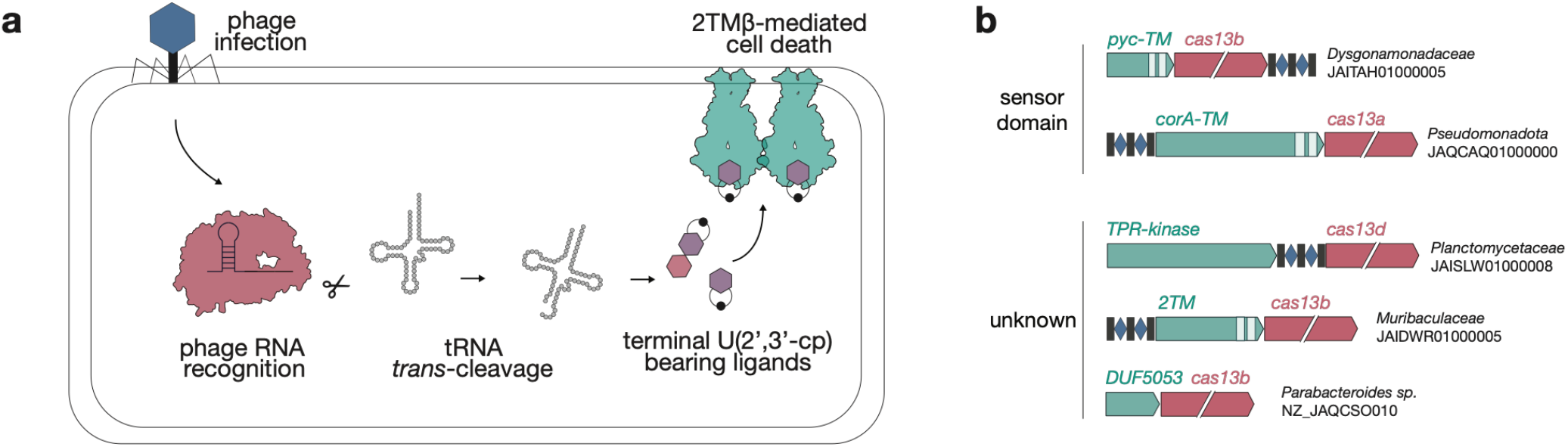
A tRNA-derived signaling model and broader Cas13–effector associations. **a**, Model for activation of the 2TMβ–Cas13b6 defense pathway. Target RNA recognition activates Cas13b6-mediated cleavage of tRNA anticodon loops, generating tRNA-derived products bearing a terminal uridine 2′,3′-cyclic-phosphate moiety. These products promote 2TMβ assembly, leading to membrane damage and loss of cell viability. **b**, Representative cas13-associated loci encoding predicted sensor-domain proteins and other proteins of uncharacterized function. Gene organization and sequence accession identifiers are shown.

Prokaryotic immune second messengers have been understood as messengers generated by enzymatic conversion of nucleotide precursors, including cUMP and cGMP produced from UTP and GTP, respectively. In CBASS, Thoeris, Pycsar and type III CRISPR immune systems, a dedicated enzyme synthesizes a cyclic nucleotide or ADP-ribose messenger by modifying cellular nucleotides, and the resulting messenger molecule triggers an effector protein to mount a response^1–3^. The pathway described here operates by the opposite logic. Rather than assembling a messenger from precursors, Cas13 cleaves tRNAs, among the most abundant RNAs in the cell, to release products bearing a terminal uridine 2′,3′-cyclic phosphate. This provides a potential route to immune signalling from an abundant, pre-existing RNA pool. Synthetic GpU(2′,3′-cp) and 2′,3′-cUMP were sufficient to promote assembly of the isolated 2TMβ β-barrel (**Fig. 3d,e**). Whether *2TMβ-cas13b6* systems also read other tRNA fragments as signals remains unknown, and the breadth of effector-gene association we observe raises the possibility of substantial ligand diversity. Our genomic survey reveals that signal sensor-effectors such as Pyc-TM, a 3′,5′-cCMP 3′,5′-cUMP sensor-effector from the Pycsar system, recur across numerous Cas13 loci, indicating repeated exaptation of these modules from other signaling pathways^8^ **(Fig. 5b)**. The logic of these findings prompts a broader reconsideration of tRNA cleavage in bacterial immunity. Targeting tRNA is a recurrent antiphage strategy, used not only by CRISPR-Cas nucleases such as Cas13^17^ and Cas12a3^21^, but also by non-CRISPR-Cas effectors such as the PrrC anticodon nuclease, the PARIS effector AriB, Retron-Eco7, RemAIN, and ApeA^22–25^. These activities are currently interpreted as terminal, with depletion of a tRNA species arresting translation to abort infection, but our findings raise the possibility that in some systems, the cleaved tRNA is not merely a depleted substrate but the input to a downstream signaling pathway.

tRNA modifications could add a further dimension to this signaling chemistry. In *E. coli* His and Asp tRNAs, G34 is a site of queuosine (Q) modification; Asp tRNA can additionally carry glutamyl-queuosine ^26,27^. Cleavage at the corresponding bonds in Q-modified substrates would therefore yield modified dinucleotides such as QpU(2′,3′-cp), making these plausible cellular counterparts of the GpU(2′,3′-cp) identified from unmodified transcripts. Our reconstitution establishes that Q is not required for cleavage of these substrates or for ligand-induced assembly *in vitro*, though whether queuosine is required for physiological signaling, or instead tunes product formation or effector activation, remains unanswered. More broadly, tRNA modifications could expand the chemical repertoire available to immune signaling beyond that specified by the four canonical ribonucleotides.

The use of tRNA fragments as signals connects bacterial immunity to a growing body of work in eukaryotes, where tRNA cleavage products act as regulatory molecules that orchestrate various cellular processes ^28^. That both prokaryotes and eukaryotes independently couple tRNA cleavage to a downstream response suggests that the cell repeatedly reads tRNA cleavage as a signal, a logic that exploits tRNA’s abundance and its position at the center of protein synthesis. Beyond its role in defense, the Cas13b6 discovered here is an unusual enzyme that cleaves tRNAs in a programmable, guide-directed manner while apparently sparing the activating transcript and the broader transcriptome. This selectivity could make it a powerful tool for studying tRNA fragmentation and the downstream pathways it controls, particularly in heterologous eukaryotic hosts in which generating specific tRNA fragments on demand would be otherwise difficult. Because specific tRNA fragments can themselves be beneficial, conferring cellular protection and even slowing aging, the ability to generate a chosen fragment on demand may also open a route to harnessing these pathways therapeutically^29,30^.

## Acknowledgements

We thank members of the Doudna and Knott laboratories for technical advice, insightful discussions, and feedback on the manuscript. We are grateful to K. Lucas for outstanding operational and administrative leadership in support of the Doudna laboratory’s research community, and to K. Zhou and J. Ye for technical support in the Doudna laboratory. We also thank David Savage for providing access to the fluorescence microscope. Mass spectra of Cas13-derived tRNA products were acquired with assistance from Dr. Tung Le in the Schepartz laboratory using an ACQUITY UPLC I-Class PLUS system coupled to a Xevo G2-XS ToF mass spectrometer (Waters). The instrument was acquired with support from the NSF Center for Genetically Encoded Materials (CHE-2002182). We thank M. West at the UC Berkeley Cell and Tissue Analysis Facility (RRID: SCR_023741) for assistance with mass photometry experiments. We acknowledge support from the following core facilities: UCSF Quantitative Metabolite Analysis Center (QMAC) partially funded by the Benioff Center for Microbiome Medicine (BCMM). We thank Undine Manzau at Biolog Life Science Institute GmbH & Co. KG for technical assistance with the synthesis and purification of nucleotide ligands. Thank you to Dr. Danielle Jorgens and Reena Zalpuri at the University of California Berkeley Electron Microscope Laboratory for advice and assistance in experiment planning, electron microscopy sample preparation, and data collection. We thank M. J. Al-Shimary for providing phages T4, T5, T7, and EdH4, and L. Marraffini for providing λvir. We also acknowledge the Innovative Genomics Institute for research support.

## Funding

This work was supported by the Howard Hughes Medical Institute, a grant from the National Science Foundation (NSF 2334028), and Snow Medical Research Foundation (SMRF2021-276). L.E.V.A. was supported by the Human Frontier Science Program (HFSP). P.H.Y. was supported by an NSF Graduate Fellowship. S.C.L was supported by the Pew Charitable Trusts and a University of California Chancellor’s Postdoctoral Fellowship. Y.S. is a Don Brown Awardee of the Life Sciences Research Foundation. G.J.K. was supported by the Snow Medical Research Foundation (SMRF2021-276). F. S. is the head of research and development of BIOLOG Life Science Institute, which sells cyclic nucleotide analogues for research purposes. NIH R35GM133462 (MRO).

## Author contributions

L.E.V.-A., P.H.Y., G.J.K. and J.A.D. conceived the study. L.E.V.-A. and P.H.Y. led the experimental design and execution of the project. P.H.Y. and S.C.L. performed cloning, bacterial genetics, phage defense assays and phageless activation assays. L.E.V.-A., P.H.Y. and K.L. performed bioinformatic analyses and interpreted comparative genomic data, with G.B. and G.J.K contributing the initial Cas13 database. L.E.V.-A. and P.H.Y. led the biochemical work, including protein purification and *in vitro* biochemical assays, with additional biochemical experiments performed by M.M.U-R, I.E.-G., and Z.T.Z.; C.T. performed collateral-cleavage assays. L.E.V.-A. and E.G.A. performed microscopy experiments. S.C.L., L.E.V.-A., M.K. and M.R.O. performed the initial cell-viability and intracellular ATP assays. Y.S. and L.E.V.-A. performed and analyzed tRNAs-cleavage and LC–MS data. D.K. synthesized GpUp. F.S. designed and supervised nucleotide synthesis. C.L. performed polysome profiling and northern blots. E.E.D. contributed expertise on 2TMβ biology and provided assistance in biochemical experiments. J.H.D.C. contributed expertise, resources and interpretation for RNA biochemistry results. J.A.D. and G.J.K. supervised the project and acquired funding. L.E.V.-A., P.H.Y., G.J.K and J.A.D. wrote the manuscript with input from all authors. All authors reviewed and approved the manuscript.

## Competing interest statement

J.A.D. is a co-founder of Caribou Biosciences, Editas Medicine, Intellia Therapeutics, Scribe Therapeutics, Mammoth Biosciences, Evercrisp and Azalea; a scientific advisory board member of Caribou Biosciences, Scribe Therapeutics, Mammoth and Inari; and a Director at Johnson & Johnson, Tempus and Altos Labs. The regents of the University of California have patents issued for CRISPR technologies on which G.J.K. is listed as an inventor. M.R.O. is an inventor on patent applications related to CRISPR-Cas systems and uses thereof. M.R.O. is a member of the scientific advisory boards for Dahlia Biosciences and an equity holder in Dahlia Biosciences. D.K. and F.S. are employed at Biolog Life Science Institute GmbH & Co. KG, which sells nucleotide analogues. The other authors declare no competing interests.

## Data Availability

RNA-sequencing data will be deposited in NCBI GEO, and plasmids will be made available through Addgene. Raw microscopy images, LC–MS and LC–MS/MS data, mass-photometry data and associated source data will be deposited in Zenodo. All datasets and plasmids will be publicly available upon publication.

## Materials and Methods

### Bioinformatic identification and classification of *2TMβ–cas13* loci

To identify candidate hybrid type VI signaling loci, we searched publicly available prokaryotic genomes from NCBI (272.0) for compact two-transmembrane β-barrel proteins (2TMβ) and then examined their surrounding gene neighborhoods for defense-associated genes, including Cas13 family nucleases using a custom python script^31^. For each 2TMβ hit, genes within a defined genomic window were extracted and functionally annotated by domain prediction and sequence-similarity searches. Loci were classified as *cas13*-associated when a *cas13-*family gene occurred in the same predicted operon or within a short neighborhood distance of the *2TMβ* gene.

*2TMβ-cas13* loci were manually curated into shared, tandem, or solo architectures on the basis of the presence or absence of additional signaling genes, including candidate cyclases and CBASS-related modules. Redundant loci were collapsed prior to phylogenetic analysis. Protein multiple-sequence alignments were generated using MAFFT^32^ separately for the 2TMβ proteins and the associated Cas13 proteins, and trees were inferred using IQ-TREE^33^ and ModelFinder from the aligned sequences to assess whether the *2TMβ–cas13* association was restricted to a single lineage or had arisen repeatedly across different clades. Representative loci shown in the main text were selected to illustrate the major architecture classes and the broader recurrence of 2TMβ across CBASS-, Panoptes-, Thoeris-, and type VI-associated neighborhoods.

### Bacterial strains and culturing

NEB 10-beta *E. coli* (New England Biolabs), 10-beta *E. coli* Mix & Go! Competent Cells (Zymo), Mach1 *E. coli* (Invitrogen), BL21(DE3) *E. coli* (New England Biolabs), *E. coli* BW25113 ^34^ and *E. coli* MG1655^35^ were used for plasmid cloning and all experiments in this study, as described in the following sections **(Supplementary Tables 1-2)**. *E. coli* strains were grown in LB broth at 37°C, 250 RPM or on 1.5% agar plates, with appropriate inducers and antibiotics. Inducers and antibiotics were used at the following working concentrations: 0.05% L-Arabinose (Sigma), 100 nM Crystal Violet (CV, Sigma), 0.8 mM Isopropyl β-D-1-thiogalactopyranoside (IPTG, Sigma), 50 nM Anhydrotetracycline Hydrochloride (aTc, Thermo Scientific Chemicals) 35 µg/mL Kanamycin (Sigma), 20 µg/mL Chloramphenicol (Sigma), and 100 µg/mL Ampicillin or Carbenicillin (Sigma).

### Cloning and plasmid construction

All plasmids were assembled by Gibson Assembly, Golden Gate Assembly, or KLD/primer-extension cloning. PCR products were generated with KAPA HiFi (Roche), PrimeStar GXL (Takara) or Q5 Hot Start High-Fidelity DNA Polymerase (New England Biolabs) PCR mixes, using IDT-synthesised primers. Final constructs were sequence-verified by Plasmidsaurus or Quintara Biosciences. Four expression systems were used throughout this study. The pJEX-SC101 vector encodes a crystal violet inducible pJEX promoter. The pBAD-ColE1 vector encodes an arabinose-inducible pBAD promoter. The pTet-p15A vector encodes an aTc-inducible pTet promoter, as well as a constitutive J23119 promoter for CRISPR guide RNA expression. The pT7-SC101 encodes an IPTG-inducible T7 promoter and a N-terminus 10xHis–Maltose Binding Protein (MBP)–TEV cleavage site tags.

To study the antiphage activity of the *Chitinophaga sp* (Cas13b6) Type VI CRISPR system, the entire locus, including the *2TMβ-cas13b6* operon and a minimal CRISPR array (Repeat + 2xBbsI sites + Repeat), was synthesised as gene fragments (Twist) and cloned into the pJEX-SC101 vector. This parent plasmid was used to generate two types of variants: modifications of the CRISPR array by Golden Gate Assembly yielded reprogrammed crRNAs for targeting of various RNAs (phage and plasmid-derived); modifications to the *2TMβ-cas13b6* operon, generated by KLD/primer-extension cloning, yielded a panel of catalytic and deletion mutants in Cas13 and 2TMβ that served to dissect the role of each of these components in the observed antiviral activity.

To study the antiphage activity of the *Chryseobacterium sp* (Cas13b7) Type VI CRISPR system, the entire locus, including the *2TMβ-cas13b7* operon and a minimal CRISPR array (Repeat + 2x BsaI sites + Repeat), was *E. coli*-codon optimized, synthesised as gene fragments (Twist) and cloned into the pTet-p15A vector: the *2TMβ-cas13b7* operon was thus placed under control of the pTet promoter, while the minimal CRISPR array was constitutively expressed by the J23119 promoter. Two types of plasmid variants were generated, as described above for the *Chitinophaga sp*. orthologue. For phage-less activation assays, a target plasmid was generated by cloning the sfGFP CDS into the pBAD-ColE1 vector. Accordingly, the wild-type *2TMβ-cas13b6* CRISPR system and mutants thereof (Δ2TMβ, Cas13b6 dHEPN1) were reprogrammed to encode a sfGFP-targeting spacer.

### Phage propagation and titration

Phages were propagated on *E. coli* BW25113 in LB at 37°C, using the webbed plate lysis method. Cleared and filtered lysates were stored at 4ºC, with their titres determined via plaque assays on empty vector-carrying *E. coli* BW25113.

### Plaque assays

Plaque assays were performed using the double agar overlay method. 100 µL of saturated overnight cultures carrying either a wild-type type VI CRISPR system or mutants thereof (as described in the main text) were added to 5 mL of molten LB agar (0.7% agar, ~50°C; henceforth, “top agar”) supplemented with antibiotics, and optionally inducer.

For plaque assays that assessed the antiviral activity of the *Chitinophaga 2TMβ-cas13b6* operon against a panel of lytic phages, top agar was supplemented with crystal violet at a final concentration of 100 nM to promote expression of the targeting complex. For plaque assays that assessed the antiviral activity of the *2TMβ-cas13b7* operon against λvir, top agar was supplemented with aTc at a final concentration of 50 nM to promote expression of the targeting complex. The supplemented top agar was poured over 1.5% LB agar plates with antibiotics, and was allowed to cool to room temperature and solidify. Tenfold phage dilutions were prepared in SM buffer (Teknova), then 2.5 µL of this dilution series was spotted onto the top agar. After drying under sterile conditions, plates were incubated at 37ºC overnight. Plaque assays were performed in biological triplicate.

### Liquid infection assays

Saturated overnight cultures carrying either a wild-type, phage-targeting *Chitinophaga 2TMβ-cas13b6* operon or mutants thereof (as described in the main text), were seeded in a microplate (Corning) at ~8e6 CFU per well in 200 µL of LB supplemented with carbenicillin and 100nM crystal violet. Tenfold dilutions of phage T7 in SM buffer were added to cells, and growth was monitored by measurement of OD600 every 5 minutes while shaking at 800 RPM (double orbital) at 37ºC overnight on a 96-well plate reader (Biotek Cytation 5).

### Phage-less activation assays

The 2TMβ-cas13 plasmids and target GFP plasmids were introduced into *E. coli* NEB 10-beta strains (New England Biolabs) via electroporation. Approximately 25 ng of each plasmid was used in 0.1 mm cuvettes on a Micropulser device (Bio-Rad). Transformed cells were recovered for 1h, and serially diluted and plated on LB-agar containing 100 µg/mL ampicillin (Sigma), 35 µg/mL kanamycin (Sigma), and optionally 100nM crystal violet (CV, Sigma). Fluorescence intensity and cell viability were measured after overnight incubation at 37ºC to assess the depletion of the GFP fluorescence and CRISPR-induced cytotoxicity.

### Expression and purification of Cas13b6 proteins

Cas13b6 constructs encoding wild-type (WT), H104A, dHEPN1 and dHEPN2 proteins were expressed in *Escherichia coli* Rosetta (DE3) pLysS cells carrying pCas13b6, which encodes an N-terminal His–TEV fusion tag. For each preparation, four 800-ml cultures were grown in 2YT medium supplemented with 100 µg/mL ampicillin (Sigma) at 37 °C to an OD600 of 0.4–0.6. Protein expression was induced with 0.5-0.8 mM IPTG, and cultures were incubated overnight at 16 °C with shaking at 160 rpm. Cells were harvested by centrifugation at 4,000g for 15 min at 4 °C, resuspended in lysis buffer consisting of 50 mM Tris-HCl pH 7.8, 500 mM NaCl, 1 mM TCEP, 10% glycerol, 0.5 mM PMSF and EDTA-free protease inhibitors, and lysed by sonication. Lysates were clarified by centrifugation at 30,000 g, 15min and 4°C, and the supernatant was applied to Ni-NTA resin equilibrated in wash buffer consisting of 50 mM Tris-HCl pH 7.8, 500 mM NaCl, 1 mM TCEP, 10% glycerol and 10 mM imidazole. The NTA resin was washed sequentially with buffers containing 10 mM and 20 mM imidazole, and bound protein was eluted with buffer containing 50 mM Tris-HCl pH 7.8, 500 mM NaCl, 1 mM TCEP, 10% glycerol and 300 mM imidazole. Eluted protein was incubated overnight at 4 °C with TEV protease while dialysing against buffer containing 50 mM HEPES pH 7.8, 250 mM NaCl, 1 mM TCEP and 10% glycerol to remove the affinity tag and reduce imidazole and salt concentrations before ion-exchange chromatography. The dialyzed sample was loaded onto a HiTrap column, Cytiva equilibrated in ion-exchange buffer A consisting of 50 mM HEPES pH 7.8, 200 mM KCl, 1 mM TCEP and 10% glycerol. Protein was eluted using a linear gradient to ion-exchange buffer B consisting of 50 mM HEPES pH 7.8, 1 M KCl, 1 mM TCEP and 10% glycerol. Peak fractions containing Cas13b6 were pooled, concentrated and further purified by size-exclusion chromatography using a Superdex 200 column (Cytiva) equilibrated in gel-filtration buffer consisting of 20 mM HEPES pH 7.8, 400 mM KCl, 1 mM TCEP and 10% glycerol. Peak fractions were pooled, concentrated to 30–50 µM, flash-frozen in aliquots and stored at −80 °C.

### *In vitro* transcription of tRNAs

DNA templates encoding tRNA^His^_GUG_, tRNA_Asp_ ^GUC^, and tRNA^Val^_GAC_ were obtained from Integrated DNA Technologies as Ultramer oligonucleotides **(Supplementary Table 3)**. Transcription templates were amplified by PCR using the forward and reverse primers listed in **(Supplementary Table 3)**; each reverse primer carried a 2′-O-methylated residue at the 5′ terminus to suppress non-templated nucleotide addition at the 3′ end of the transcript. Reactions contained 5 µg of Ultramer template, 0.5 µM each primer, and Q5 Hot Start High-Fidelity 2X Master Mix (New England Biolabs) in a 200 µL reaction. Amplification was confirmed by 1% agarose gel electrophoresis, and the unpurified reaction was used directly as template for transcription.

Transcription reactions (1 mL) were assembled in 30 mM Tris-HCl pH 7.5, 10 mM DTT, 0.1% spermidine, 0.1% Triton X-100 supplemented with 60 mM MgCl_2_, 6 mM of each NTP, T7 RNA polymerase (purified in-house), and 20% (v/v) of the PCR reaction. After incubation at 37 °C for 8 hr, the insoluble inorganic pyrophosphate was pelleted by centrifugation at 20,000 x g for 1 minute, and then the RNA-containing supernatant was recovered directly by ethanol precipitation and resolved on a 15% polyacrylamide gel containing 7 M urea in 1X TBE. The band corresponding to full-length tRNA was located by UV shadowing and excised, and RNA was eluted from the crushed slice in nuclease-free water with 20 mM NaOAc pH 5.2 and 1 mM EDTA overnight at 4 °C. Gel fragments were removed by vacuum filtration through a 0.2 µm filter unit (Thermo Scientific), and the filtrate was concentrated and exchanged into nuclease-free water with 3 mM NaOAc pH 5.2 and 0.5 mM EDTA using 3 kDa MWCO Amicon Ultra centrifugal filters (Millipore). Purified IVT tRNA was stored at −80 °C.

### Fluorogenic RNA reporter cleavage assays

Cas13-mediated collateral RNA cleavage activity was measured using a fluorescent poly(U) (6FAM/UUUUU/IowaBlackQ) RNA reporter. Prior to reaction assembly, crRNA and activator RNA were annealed at 65°C for 3 min and immediately cooled on ice for 10 min. Cas13 ribonucleoprotein (RNP) complexes were assembled by incubating Cas13 protein with crRNA at a final concentration of 300 nM each for 1 h at 4 °C for Cas13b6, or for 15 min at room temperature for LbuCas13a and RfxCas13d. Cleavage reactions were performed in a final volume of 20 µl. Cas13b6 reactions were carried out in 20 mM HEPES– KOH (pH 7.5), 200 mM KCl, 2 mM MgCl_2_, 1 mM TCEP and 5% (v/v) glycerol. LbuCas13a and RfxCas13d reactions were performed in 20 mM HEPES, 50 mM KCl, 5 mM MgCl_2_, 5% (v/v) glycerol and 0.01% IGEPAL, prepared by dilution from a 5X stock solution. Reactions were initiated by adding a solution containing the fluorescent reporter (600 nM final concentration) and activator RNA at the indicated concentrations to the preassembled RNP complexes. Fluorescence was monitored at 37 °C every 30 s for 1 h. Each condition was analyzed in three technical replicates.

### In vitro cleavage of E. coli tRNAs

Purified Cas13b6 variants were tested for activated RNA cleavage activity *in vitro* using *E. coli* tRNAs as substrates. Reactions were assembled in 50 µL with 1X Cas13 reaction buffer consisting of 20 mM HEPES, pH 7.5, 200 mM KCl, 2 mM MgCl_2_, 1 mM TCEP, 5% glycerol and 0.005% IGEPAL. Each reaction contained 200 nM Cas13 protein, 240 nM guide RNA, 300 nM activator RNA and 5 µg RNA substrate. Cas13 proteins tested included wild-type Cas13b6, H104A, and catalytic-site HEPN mutants, including dHEPN controls. Cas13–guide RNA complexes were preassembled for 10–15 min at 37°C or on ice for 60 min, activated by addition of target/activator RNA for 5–10 min, and reactions were initiated by addition of RNA substrate. Reactions were incubated at 37°C and sampled at the indicated time points or after 30–60 min for endpoint assays. Samples were quenched in denaturing RNA loading dye and resolved on 15% TBE–urea polyacrylamide gels. RNA was visualized by SYBR gold and imaged using Typhoon. Cleavage was monitored by depletion of intact tRNA-sized RNA species and accumulation of lower-molecular-weight RNA fragments.

### Time-course analysis of target-RNA and tRNA cleavage

Time-course cleavage assays contained 200 nM WT Cas13, 300 nM GFP gRNA, 300 nM 60-nt FAM-labelled GFP target RNA and 0.5 mg ml^−1^ purified E. coli tRNA in 1× Cas13 reaction buffer (20 mM HEPES, pH 7.5, 200 mM KCl, 2 mM MgCl_2_, 1 mM TCEP, 5% glycerol and 0.005% IGEPAL CA-630). Reactions were initiated by addition of the 60-nt target RNA. An aliquot was removed and quenched immediately after target addition (0 min), and additional aliquots were collected after 1, 2, 5, 10, 30 and 60 min at 37 °C. Reactions were quenched with an equal volume of 2× denaturing RNA loading dye containing EDTA, heated for 5 min at 75 °C and resolved by 15% PAGE containing 7 M urea. FAM fluorescence was used to monitor the target RNA and its cleavage products, and the same gel was stained with SYBR Gold to visualize total RNA and tRNA cleavage products.

### Cleavage of synthetic tRNA anticodon stem–loops

Synthetic 21-nt RNA oligonucleotides carrying a 5′ Cy5 label were used to examine cleavage of isolated anticodon stem-loops (ASLs) from tRNA^Val-GAC^ and tRNA^His-GUG^. Each construct comprised the reference-derived tRNA segment corresponding to positions 27–43, extended by 5′-GG and 3′-CC to provide two additional stem base pairs. Val ASLs containing U33A or U33A/G34U substitutions were tested alongside the parental sequence. Residues were numbered relative to anticodon positions 34–36. Oligonucleotide sequences are provided in Supplementary Table 3.

Cleavage reactions were performed in a final volume of 20 µl in 1× Cas13 reaction buffer, as described above, with final concentrations of 200 nM WT Cas13b6 or H104A, 300 nM GFP-targeting guide RNA and 200 nM Cy5-labelled ASL substrate. Cas13–guide RNA complexes were preassembled for 15 min at 37 °C. FAM-labelled 60-nt GFP activator RNA was then added to a final concentration of 200 nM, followed by incubation for 5 min at 37 °C. For non-activated controls, activator RNA was replaced with an equal volume of nuclease-free water. Cleavage was initiated by addition of the ASL substrate, and reactions were incubated for 60 min at 37 °C. Reactions were quenched with an equal volume of 2× denaturing RNA loading dye containing EDTA and resolved by 20% TBE–urea PAGE. Intact ASLs and cleavage products retaining the 5′ label were visualized by Cy5 fluorescence. Reactions were performed in duplicate.

### Small RNA-seq and mapping of Cas13 cleavage sites

For end mapping, the 1 h cleavage reaction of *E. coli* total RNA was purified with SPRI beads and oxidized with 50 mM sodium periodate in 0.1 M sodium acetate (pH 5.2) for 30 min at 25 °C in the dark. The reaction was quenched with glycerol and purified by SPRI. Oxidized RNA was treated with T4 polynucleotide kinase in 1X T4 PNK buffer, first without ATP to convert 2′,3′-cyclic phosphate termini to 3′-OH and then with ATP to 5′-phosphorylate. Small-RNA sequencing libraries were prepared with the Collibri small-RNA-seq library preparation following manufacturer protocol. Libraries were size-selected by gel extraction from 4% E-Gel EX Agarose Gels for inserts of 30–300 nt and sequenced with paired-end 150 bp reads via Illumina Next-Seq 1000. Sequencing reads were imported into Geneious and read pairs merged with BBMerge. Merged reads were mapped to the *E. coli* str. K-12 substrain DH10B reference complete genome (GenBank CP000948.1^36^) with the Geneious mapper at low sensitivity. Reads were mapped in parallel to the whole genome, to ribosomal RNA, and to tRNA sequences to identify the most abundant mapped species. Manual inspection showed altered coverage across tRNA loci, and read 5′ and 3′ end positions were then plotted per nucleotide to localize cleavage termini.

SuperasIN and 8 uL of 180mM Tris pH9 for deacylation at 37C for 30 min. Next 3′ end repair was done with addition of 5 µL of T4 PNK mix (200 mM Tris-HCl pH 6.8, 40 mM MgCl 2, 4 U/µL T4 PNK, from New England Biolabs) for 30min at 37C, followed by deactivation at 65C for 10 min. Next Hairpin adaptors barcoded oligos was added – 2uL of 20uM to each sample, followed by addition of 28uL of ligation mix: 1 U/µL T4 RNA ligase I (NEB), 1x NEB T4 RNA ligase I buffer, 15% PEG 8000, 50 µM ATP, 1 mM hexamine cobalt chloride, and 5% DMSO. Ligation was done for 12 hours at 16C. Samples were then bound to streptavidin beads, pooled, and dephosphorylated. Then cDNA was synthesized with SSIV at 55C for 10 min followed by 35 C overnight. Next RNA was digested with RNaseH, empty adaptor was inactivated by periodate oxidation, and a second adaptor was ligated at room temperature overnight. cDNA was boiled and moved to a new tube to released form streptavidin beads then used as template for 12 cycles of PCR with Q5 polymerase and indexing primers. Amplified libraries were gel cut between 150and 350bp of size, purified, and used for sequencing on an Illumina NovaSeq X plus. Bioinformatics was additionally performed by MesoRNA (Chicago, IL, USA). Reads were demultiplexed and trimmed with custom software in accordance with the MSR-seq library preparation protocol^37^. Quality control was performed with FastQC. Reads were aligned to tRNA sequences adapted from GtRNAdb^38^with bowtie2^39^.

### Northern blot analysis

Commercially purified *E. coli* tRNA (New England Biolabs, N6845S) was used as substrate for *in vitro* Cas13b6 cleavage reactions. The tRNA preparation was assessed by denaturing PAGE before use, and no additional size-based purification was performed. Cas13b6 WT, H104A or dHEPN1 was pre-incubated with GFP-targeting gRNA for 15 min at 37 °C in 1× Cas13 reaction buffer (20 mM HEPES, pH 7.5, 200 mM KCl, 2 mM MgCl_2_, 1 mM TCEP, 5% glycerol and 0.005% IGEPAL CA-630). Reactions contained 500 nM Cas13b6, 500 nM gRNA, 500 nM unlabeled 122-nt GFP activating RNA (aRNA) and 0.5 mg ml^−1^ purified *E. coli* tRNA in a final volume of 25 µl. For non-activated controls, aRNA was omitted and replaced with nuclease-free water. Following RNP assembly, aRNA was added and incubated for 5 min at 37 °C before addition of tRNA to initiate cleavage. Reactions were incubated for 1 h at 37 °C and quenched with an equal volume of 2× denaturing RNA loading dye containing EDTA.

RNA (0.3 µg per lane) was resolved on a 15% polyacrylamide, 7 M urea denaturing gel in 1× TBE at 100 V for 1.5 h. RNA was transferred to positively charged Hybond-N+ nylon membrane (GE Healthcare) by wet electrotransfer in 0.5× TBE at 100 V for 1 h at 4 °C and crosslinked to the membrane by UV irradiation at 120 mJ cm^−2^ using a Spectrolinker XL-1500 (Spectronics). Membranes were prehybridized in 5× SSC containing 1% SDS for 1 h at 42 °C and subsequently hybridized for 12 h at 42 °C in ULTRAhyb-Oligo buffer (Ambion) containing 2 nM 5′-Cy5-labelled DNA oligonucleotide probe. Sequence-specific probes complementary to the 5′ region shared by the ValV and ValW tRNA^Val^ _GAC_ isoacceptors, the isoacceptor-specific 3′ regions of ValV and ValW, and the 5′ region of tRNA^His^ _GUG_ were used to detect full-length and cleaved tRNA species. Where indicated, a 5′-Cy3-labelled probe complementary to 5S rRNA was included as a loading control. Oligonucleotide sequences are provided in Supplementary Table 4. Membranes were washed twice with 2× SSC containing 0.1% SDS for 10 min at 42 °C and imaged using a Typhoon fluorescence scanner (Cytiva) in the Cy5 and, where applicable, Cy3 channels.

### RNA extraction and gel electrophoresis

Total RNA was extracted from clarified total lysates by acid phenol-chloroform extraction followed by chloroform back-extraction and ethanol precipitation. RNA concentration was determined by A254 absorbance, and 200 ng per lane was resolved on a 1.5% native agarose gel in 1xTAE stained with SYBR Safe.

### Cas13 cleavage of IVT tRNAs and isolation of RNA products for LC– MS

Libraries were prepared by MesoRNA (Chicago, IL, USA) following the MSR-seq protocol as previously described (Watkins, 2021). Briefly, 1ug of total RNA was diluted into 6uL of water, then mixed with 1ul of For cleavage-product analysis by intact tRNA LC–MS, Cas13b6 ribonucleoprotein (RNP) complexes were assembled in 50 µl reactions containing Cas13 reaction buffer (20 mM HEPES pH 7.5, 200 mM KCl, 2 mM MgCl_2_, 1 mM TCEP, 5% glycerol and 0.005% IGEPAL), 200 nM Cas13b6 protein, 240 nM guide RNA and, where indicated, 300 nM activator RNA. RNPs were preassembled on ice for 60 min and activated for 15 min at 37 °C before addition of IVT tRNA^Ala^ to a final concentration of 1 µg µl^−1^. Reactions were incubated at 37 °C for 1 hour. For gel-based analysis, aliquots were quenched in denaturing RNA loading dye and resolved on 15% TBE–urea polyacrylamide gels. For LC– MS analysis, cleavage reactions were transferred to 30-kDa MWCO Amicon Ultra centrifugal filters to remove Cas13 RNP and other high-molecular-weight components. The filtrate, containing full-length RNA substrate and low-molecular-weight cleavage products, was then purified using a Thermo Fisher miRNA cleanup workflow according to the manufacturer’s protocol and eluted in nuclease-free water. Purified RNA was used directly for intact RNA LC–MS.

### PNK-dependent adapter ligation

Cas13-generated cleavage products from *in vitro* transcribed Val, His and Asp tRNAs were prepared and purified using the mirVana miRNA Isolation Kit (Thermo Fisher Scientific), as described for intact-RNA LC–MS, and used directly without further fractionation. Synthetic tRNA^Val^ and tRNA^His^ 5′ fragments (ChemGenes Corporation) bearing defined 2′,3′-cyclic-phosphate termini were analyzed in parallel as reference substrates. RNA and adapter sequences are provided in Supplementary Table 4. Approximately 1 µg of each purified Cas13 cleavage-product preparation or 40 pmol of synthetic RNA was incubated with 10 U T4 polynucleotide kinase (PNK; New England Biolabs, M0201) in a 10-µl reaction containing 1× T4 PNK Reaction Buffer, without added ATP. Mock-treated controls contained the same components, with PNK replaced by nuclease-free water. Samples were incubated for 30 min at 37 °C, followed by 20 min at 65 °C to inactivate PNK, and cooled on ice.

For adapter ligation, 5 µl of each PNK-treated or mock-treated sample was added to a 20-µl reaction containing 2 µl of 10× T4 RNA Ligase Reaction Buffer, 4 µl of 50% PEG 8000, 40 pmol of the 17-nt, 5′-preadenylated, 3′-blocked Universal miRNA Cloning Linker (New England Biolabs, S1315) and 200 U T4 RNA ligase 2, truncated KQ (New England Biolabs, M0373). No ATP was added. Reactions were incubated for 1 h at 25 °C. Aliquots of PNK-treated RNA without subsequent adapter ligation were retained as controls. Samples were mixed with an equal volume of 2× denaturing RNA loading dye containing EDTA, heated for 5 min at 75 °C, cooled briefly on ice and resolved on a 15% polyacrylamide gel containing 7 M urea. RNA was visualized by SYBR Gold staining. PNK-dependent adapter ligation was assessed by comparing the appearance of slower-migrating products in PNK-treated and mock-treated samples, with the corresponding synthetic cyclic-phosphate substrates and PNK-only samples serving as controls.

### Intact-RNA LC–MS and mass deconvolution

Purified tRNA cleavage products from Cas13 *in vitro* reactions were analyzed by LC-HRMS on an ACQUITY UPLC BEH C18 column (130 Å, 1.7 µm, 2.1 X 50 mm) at 60 °C using an ACQUITY UPLC I-Class PLUS system coupled to a Waters Xevo G2-XS ToF mass spectrometer operated in negative ion mode. Mobile phase A consisted of 8 mM triethylamine (TEA), 80 mM hexafluoroisopropanol (HFIP) and 5 µM EDTA in Milli-Q water. Mobile phase B consisted of 4 mM TEA, 40 mM HFIP and 5 µM EDTA in 50% Milli-Q water / 50% methanol. The flow rate was 0.3 mL min^-1^. The gradient began at 22% B, increased linearly to 40% B over 10 min, increased to 60% B over 1 min, was held at 60% B for 1 min, returned to 22% B over 0.1 min, and was re-equilibrated at 22% B for 2.9 min. Source parameters were: capillary voltage, 2,000 V; sampling cone, 40; source offset, 40; source temperature, 140 °C; desolvation temperature, 20 °C; cone gas flow, 10 L h^-1^; desolvation gas flow, 800 L h^-1^; acquisition rate, 1 spectrum s^−1^. Expected masses for full-length IVTs and predicted cleavage products were calculated *in silico* (https://www.aatbio.com/tools/calculate-RNA-molecular-weight-mw), including products bearing 5′-hydroxyl and 2′,3′-cyclic phosphate termini. Deconvoluted mass spectra were generated using MaxEnt software (Waters), and observed masses were assigned by comparison to predicted masses of full-length RNA and candidate cleavage products. This LC–MS/deconvolution workflow was adapted from the intact tRNA LC–MS protocol^40^.

### Targeted LC–MS/MS analysis of GpU(2′,3′-cp)

Cas13 cleavage reactions were performed in a total volume of 200 µl as described above and passed through 3-kDa molecular-weight-cut-off centrifugal filters. The filtrates were collected and extracted with methanol:acetonitrile:water (40:40:20, v/v/v). Extracts were dried in a SpeedVac vacuum concentrator at room temperature, and the dried residues were reconstituted in 50 µl of LC–MS-grade water. Samples were analyzed using a SCIEX ExionLC UPLC system coupled to a SCIEX 7500 triple-quadrupole mass spectrometer. A 5-µl aliquot was injected onto an Acclaim Polar Advantage II column (3-µm particle size, 2.1 × 150 mm; Thermo Scientific). Mobile phases A and B contained 3% and 97% methanol, respectively, each supplemented with 0.1% formic acid and 50 mM ammonium formate. Separation was performed at a flow rate of 200 µl min^−1^ using the following gradient: 0.2% B from 0 to 2 min, a linear increase to 49% B at 8 min, a return to 0.2% B at 8.1 min and equilibration at 0.2% B until 10 min.

Data were acquired in positive electrospray ionization mode using multiple-reaction monitoring (MRM). Source settings were GS1 = 60, GS2 = 70, CUR = 40 and CAD = 9, with a source temperature of 375 °C and an ion-spray voltage of 2,200 V. The entrance potential was 10 V, the collision-cell exit potential was 15 V, the dwell time was 10 ms and the inter-transition pause time was 5 ms. An authentic GpU(2′,3′-cp) standard, guanylyl-(3′→5′)-uridine-2′,3′-cyclic monophosphate sodium salt, was custom-synthesized by Biolog (product no. G 085-01) and used for MRM optimization and external calibration. GpU(2′,3′-cp) was monitored using the quantifier transition m/z 652 → 152 and qualifier transitions m/z 652 → 289 and 652 → 501. The corresponding collision energies were 25, 40 and 20 V, respectively (Supplementary Table 9).

A nine-level external calibration series of GpU(2′,3′-cp) was analyzed in duplicate before and after the sample sequence and fitted using a quadratic model with 1/x weighting (R^2^ = 0.9989). A pooled quality-control sample was analyzed before the sample sequence and after every ten samples. The pre-extracted Cas13 reaction samples were analyzed without an added internal standard. Data was processed using the Analytics module of SCIEX OS software (version 2.1.6.59781).

### Structure prediction for 2TMB with second messenger ligands

Structures of dimeric *Chitinophaga* 2TMB were predicted as full-length proteins using Protenix-v1^41^. Ligand bound states were predicted across five random model seeds with five samples, 10 recycles, and 200 diffusion steps (protenix_base_20250630_v1.0.0) with 2 copies of each ligand (2′,3′-cCMP, 2′,3′-cAMP, 2′,3′-cGMP, or 2′,3′-cUMP) input as a ligand SMILES defined from PubChem Compound records.

### Expression and purification of soluble MBP–β-barrel protein

Soluble 2TMβ-barrel constructs were expressed as N-terminal His_6_– MBP fusion proteins in Rosetta (DE3) pLysS. Wild-type and mutant constructs, including Y157A, Y197A, Y197F deletion mutants, were generated by site directed mutagenesis with PCR and golden gate assembly. Cells were grown at 37 °C to an OD600 of 0.4–0.6, induced with 0.8 mM IPTG and expressed overnight at 16 °C. Cell pellets were resuspended in lysis buffer containing 20 mM HEPES, pH 7.5, 500 mM NaCl, 10 mM imidazole, 1 mM TCEP, 5% glycerol and Roche protease inhibitors, and lysed by sonication. Clarified lysates were purified by Ni-NTA affinity chromatography, washed with buffer containing 40 mM imidazole and eluted with 300 mM imidazole. Eluted proteins were further purified by size-exclusion chromatography in 20 mM HEPES, pH 7.5, 300 mM KCl and 1 mM TCEP. Peak fractions were pooled, concentrated, flash-frozen and stored at −80 °C. The MBP tag was retained for downstream ligand-binding, mass-photometry and oligomerization assays.

### Mass photometry analysis of soluble β-barrel oligomerization

Mass photometry measurements were acquired on a Refeyn TwoMP mass photometer (Refeyn Ltd) using MassFerence™ P1 calibrant for calibration. Data were analyzed in MassGlass and DiscoverMP Analysis. Purified soluble MBP–β-barrel proteins were incubated in 50 µL reactions containing 1 µM protein in 20 mM HEPES, pH 7.5, 300 mM KCl and 1 mM TCEP. The agonist 2′,3′-cUMP was purchased from BIOLOG Life Science Institute as the sodium salt (Cat. No. U 004; >99% HPLC purity) and prepared as a 230 mM stock in water. Ligands were added from concentrated stocks to a final concentration of 300 µM and incubated on ice for 15–30 min. Samples were diluted 1:20 immediately before measurement by mixing 1 µL reaction with 19 µL measurement buffer, yielding final on-slide concentrations of 50 nM protein and 15 µM ligand. WT and mutant proteins were compared under matched apo and ligand-treated conditions using proteins normalized to 10 µM working stocks. Control ligands included c-triAMP, 3′,2′-cUAMP, 3′,3′-cdUMP, 2′,3′-cCMP, 2′,3′-cAMP, 2′,3′-cGMP, 2′-UMP, 3′-UMP and 5′-UMP (Supplementary Table 5). Apparent mass distributions were compared to assess ligand-dependent oligomerization.Ligand-comparison reactions contained 1 µM MBP–2TMβ β-barrel and 300 µM ligand before dilution. GpU(2′,3′-cp) titrations contained 100, 200 or 300 µM ligand at the same protein concentration. Immediately before measurement, 1 µl of reaction was mixed with 19 µl of ligand-free measurement buffer, yielding 50 nM protein and 15 µM ligand for the comparison experiments, or 5, 10 and 15 µM ligand for the titration, respectively. Apo controls were handled with the same dilution procedure. Ligand-dependent assembly was assessed from the distributions of apparent particle masses. shaking for 5 min on a 96-well plate reader (Biotek Cytation 5), after which the luminescence was read at 590/35.

### DiBAC membrane depolarization assays

Saturated overnight cultures carrying either a wild-type, GFP-targeting, *Chitinophaga 2TMβ-cas13b6* operon or mutants thereof (as described in the main text), were back-diluted to OD 0.1 and grown to OD 0.4. Then, cultures were seeded in a 96-well microplate (Corning) at a final OD of 0.1 per well in 200 µL of LB supplemented with carbenicillin, kanamycin, and optionally, 100nM crystal violet and 0.05% L-Arabinose. Growth was monitored by measurement of OD600 every 5 minutes while shaking at 800 RPM (double orbital) at 37ºC on a 96-well plate reader (Biotek Cytation 5) until the wild-type induced cultures arrested (typically, 3h post-seeding). For membrane depolarization measurements, cells were combined with LB and DiBAC_4_(3) (Invitrogen, B438) in black 96-well assay plates (Corning), yielding afinal volume of 200 µl and a final dye concentration of 0.5 µM. Plates were incubated at 37 °C with shaking for 5 min, after which fluorescence was centrifugation at 5,000xg for 10 minutes at 4°C, washed once in ice-cold polysome buffer (20 mM Tris-HCl pH 7.5, 10 mM MgCl_2_, 100 mM NH_4_Cl, 2 mM DTT) containing 100 µg/mL chloramphenicol, and resuspended in lysis buffer (polysome buffer supplemented with lysozyme to 1 mg/mL and RNase-free DNase I). Cells were lysed by incubation on ice for 15 minutes followed by three freeze-thaw cycles, after which sodium deoxycholate was added to a final concentration of 0.25% (w/v) and incubated on ice for an additional 15 minutes. Lysates were clarified by centrifugation at 20,000xg for 15 minutes at 4°C and A260 absorbance was measured to normalize input across samples. Approximately 20 A260 units of clarified lysate were layered onto 12 mL linear 10–40% sucrose gradients prepared in polysome buffer. Gradients were centrifuged in a Beckman SW41 swinging-bucket rotor at 39,000 rpm for 3 hours at 4°C and fractionated with continuous A254 monitoring using an ISCO fractionation system.

### Transmission electron microscopy

Cells were fixed with 2.5% glutaraldehyde and 2.5% paraformaldehyde in 1x phosphate buffered saline (PBS) then incubated onto poly-lysine coated 35mm MatTek glass-bottom dishes (#P35G-1.5-14 MatTek Corp., Ashland, MA, USA) for an hour to adhere cells to the coverslip surface. Cells were rinsed (3×; 5 min, Room Temperature) in 1x PBS and incubated in 1% osmium tetroxide with 1.6% potassium ferricyanide in 1x PBS for 30 minutes. Samples were rinsed (3×; 5 min, RT) in buffer and briefly washed with distilled water (1×; 1 min, RT) before sample were then subjected to an ascending ethanol gradient followed by pure ethanol. Samples were progressively infiltrated (using ethanol as the solvent) with Epon resin (EMS, Hatfield, PA, USA) and polymerized at 60 C for 24 hours. Following polymerization, the glass coverslips were removed using ultra-thin Personna razor blades (EMS, Hatfield, PA, USA). The polymerized resin from the MatTek plate was mounted on a blank resin block with cyanoacrylate glue for sectioning. Thin sections (90 nm) were cut using a Leica UC6 ultramicrotome (Leica, Wetzlar, Germany) and collected onto formvar-coated slot grids. The grids were post-stained with 2% uranyl acetate followed by Reynold’s lead citrate, for 4 min each. Sections were imaged using a FEI Tecnai 12

### Intracellular ATP production assays

Saturated overnight cultures carrying either a wild-type, GFP-targeting, were back-diluted to OD 0.1 and grown to OD 0.4. Then, cultures were seeded in a 96-well microplate (Corning) at a final OD of 0.1 per well in 200 µL of LB supplemented with carbenicillin, kanamycin, and optionally, 100nM crystal violet and 0.05% L-Arabinose. At this point, the plates were sampled for initial (t=0) ATP measurements, described below. Growth was monitored by measurement of OD600 every 5 minutes while shaking at 800 RPM (double orbital) at 37ºC on a 96-well plate reader (Biotek Cytation 5) until the wild-type induced cultures arrested (typically, 3h post-seeding). Then, cultures were sampled for ATP measurement, as follows: 8 uL of cells were diluted in 192 uL of LB, and 50 uL of this mixture was added to 50 uL of BacTiter-Glo Reagent (Promega) in a black 96-well assay plate (Corning). The plate was incubated at 37ºC with measured using a Cytation 5 plate reader (BioTek) at excitation and emission wavelengths of 493 and 516 nm, respectively.

### Polysome profiling

Saturated overnight cultures carrying either a wild-type, GFP-targeting *Chitinophaga 2TMβ-cas13b6* operon or mutants thereof (as described in the main text), were back-diluted to OD 0.1 and grown to OD 0.4 in 200mL LB supplemented with carbenicillin and kanamycin. Then, cultures were back-diluted to a final OD of 0.1 in flasks containing 200mL of LB supplemented with carbenicillin, kanamycin, 100nM crystal violet and 0.05% L-Arabinose. In parallel, an aliquot of these cultures was seeded in a 96-well microplate (Corning) in 200 µL triplicates to monitor cell growth by measurement of OD600 every 5 minutes while shaking at 800 RPM (double orbital) at 37ºC on a 96-well plate reader (Biotek Cytation 5). Growth was monitored until the wild-type induced cultures were arrested (typically, 3h post-seeding). Then, translation was arrested by addition of chloramphenicol to a final concentration of 100 µg/mL directly to the culture for 5 minutes, after which cultures were immediately placed on ice. Cells were harvested by 120kV TEM (FEI, Hillsboro, OR, USA) and data recorded using a Gatan Rio 16 CMOS camera with Gatan Microscopy Suite software (Gatan Inc., Pleasanton, CA, USA).

### Fluorescence microscopy

Saturated overnight cultures carrying either a wild-type, GFP-targeting*Chitinophaga 2TMβ-cas13b6* operon or mutants thereof (as described in the main text), were back-diluted to OD 0.1 and grown to OD 0.4. Then, cultures were seeded in a 96-well microplate (Corning) at a final OD600 of 0.1 per well in 200 µL of LB supplemented with 100 µg/mL carbenicillin (Sigma), 20 µg/mL kanamycin (Sigma), and optionally, 100nM crystal violet and 0.05% L-Arabinose. Growth was monitored by measurement of OD600 every 5 minutes while shaking at 800 RPM (double orbital) at 37ºC on a 96-well plate reader (Biotek Cytation 5) until the wild-type induced cultures arrested (typically, 3h post-seeding). Then, cultures were harvested from the 96-well plate by gentle centrifugation (~2000g for 3min) in microcentrifuge tubes and stained in the dark for thirty minutes at 37°C. Membrane-compromised cells were labelled with SYTOX Green at a final concentration of 1–5 µM, membranes were stained with FM4-64FX or FM4-64 at 5–10 µg ml−1, and DNA was labelled with DAPI at 5 µg ml−1 where indicated. After staining, cells were washed or directly resuspended in PBS and 5 µL were spotted on agarose imaging pads (100-25% LB + 1.5% agarose). Following 1 min of drying, the pad was covered by a small coverslip (Fisher Scientific) and imaged immediately using an Axio Observer microscope (Zeiss). Matched samples were imaged using identical acquisition settings. Images were collected in at least biological duplicate.

### Identification and annotation of ***cas13-***associated genes

Subtype-specific *cas13* HMM profiles were curated and used to conduct HMMER^42^ searches against the NCBI non-redundant protein database^43^. Hit FASTA files were split by subtype and HEPN half for Cas13a, Cas13b, Cas13bt, Cas13c, Cas13d, and Cas13e. Across these files, 2,919 hit entries were deduplicated to 1,402 unique protein accessions, and 1,393 accessions were retained for genomic neighborhood retrieval.

For each retained Cas13 protein, genomic context was retrieved from NCBI IPG/GenBank using a +/-20 kb window around the focal protein. Retrieved loci were trimmed to +/-10 kb around the gene of interest, yielding 1,340 *cas13-*centered loci. Whole-locus nucleotide sequences were clustered with MMseqs2 using ‘--min-seq-id 0.5’, ‘-c 0.5’, ‘--dbtype 2’, and ‘--cluster-mode 2’, and one representative locus was retained per cluster. This produced 880 representative loci for downstream neighborhood analysis.

Non-Cas13 neighboring proteins were parsed from the 880 representative loci and clustered with the neighborhood-analysis scripts. Briefly, neighbor proteins were converted to FASTA, indexed with ‘mmseqs createdb’, and clustered with ‘mmseqs cluster’ using connected-component clustering (‘--cluster-mode 1’). Cluster assignments were exported with ‘mmseqs createtsv’, and cluster MSAs were generated with ‘mmseqs result2msa’ for HH-suite comparison. The resulting MMseqs2 protein clusters were compared with HHblits and merged into broader superclusters using probability >= 90%, query coverage >= 50%, and target coverage >= 50%. This produced 4,242 HHblits superclusters containing 8,432 unique clustered neighbor proteins.

All 8,432 clustered proteins were annotated with DeepTMHMM to identify predicted transmembrane topology and helix boundaries. DeepTMHMM calls were obtained for all proteins, yielding 1,419 TM-positive proteins distributed across 746 superclusters. Pfam annotation was performed with two complementary approaches. First, all clustered proteins were searched against Pfam-A with HMMER ‘hmmscan’ using Pfam gathering thresholds. Second, multi-sequence TM-positive clusters were aligned with MAFFT L-INS-i, converted to HH-suite HHM profiles, and searched against the HH-suite Pfam database with HHsearch. The final key TM/Pfam table integrated DeepTMHMM topology, sequence-level HMMER/Pfam calls, and cluster-level HHsearch/Pfam calls. Of 746 TM-positive superclusters, 514 had Pfam support from at least one annotation layer.

