## Supplementary Data for "tRNA-derived second messengers confer bacterial immunity"

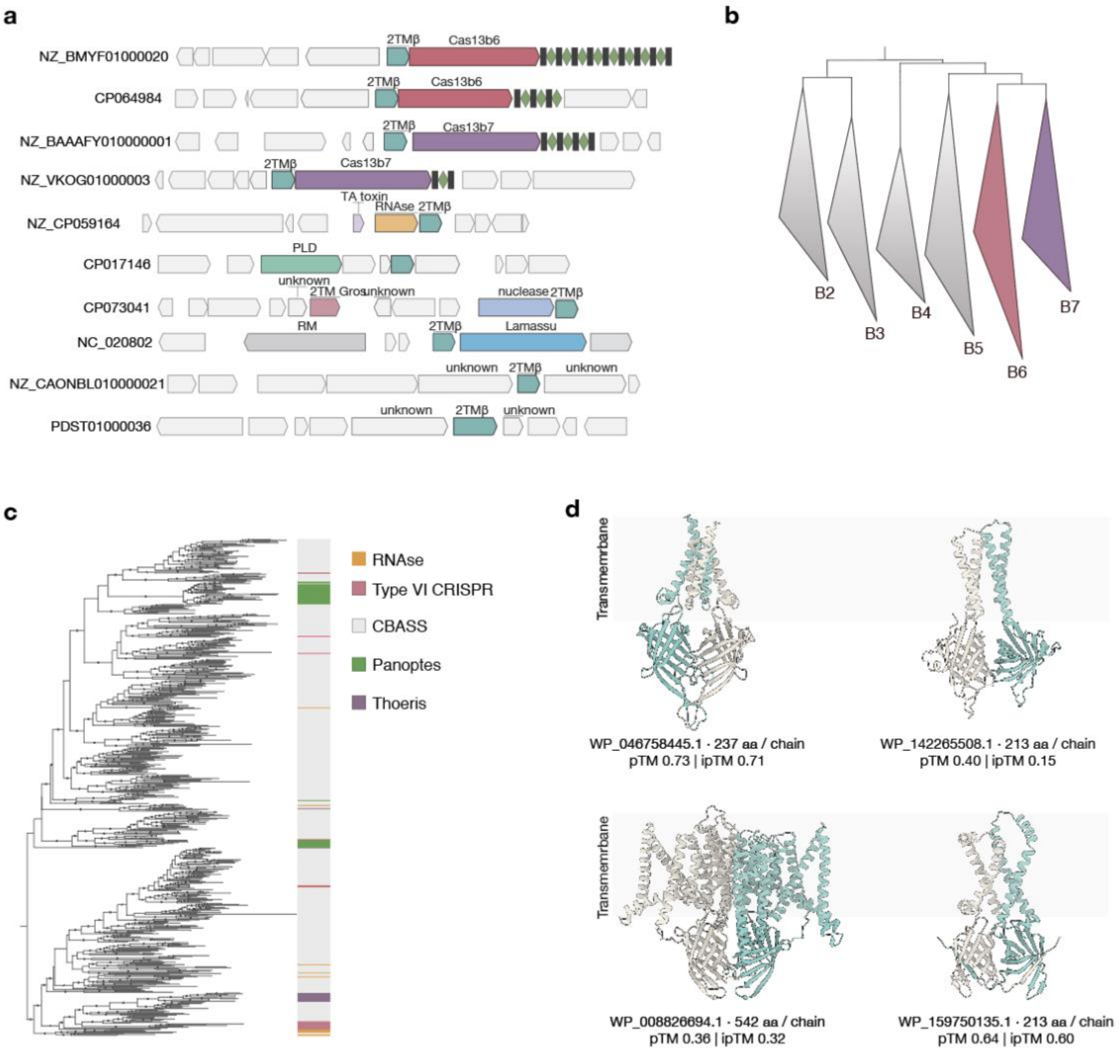


**Extended Data Fig. 1. 2TMβ proteins occur in multiple immune-gene contexts and show diverse predicted barrel architectures. a,** Representative genomic neighborhoods illustrating recurrent associations between genes encoding predicted 2TMβ proteins and diverse defense-associated genes. Gene arrows indicate coding sequences and orientation; grey arrows denote neighboring genes without assigned defense annotations**. b,** Phylogenetic placement of the *cas13b6* and *cas13b7* clades within the broader Cas13b family. **c**, Maximum-likelihood phylogeny of predicted 2TMβ proteins. **d,** Predicted structures of representative novel 2TMβ proteins from distinct genomic contexts.

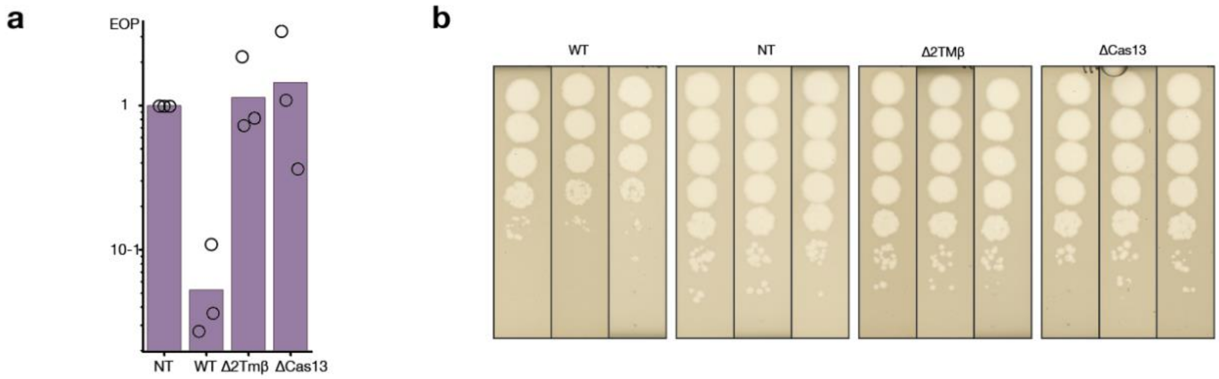


**Extended Data Fig. 2. *2TMβ-cas13b7* confers phage defense. a,** Efficiency of plating (EOP) for the *Chryseobacterium 2TMβ-cas13b7* system and the indicated deletion variants in *E. coli*. Points indicate independent biological replicates and bar heights indicate the mean. **b,** λvir spot-dilution assays corresponding to **a**. In both **a,** and **b**, Δ indicates gene deletion; WT, targeting spacer; NT, non-targeting spacer.

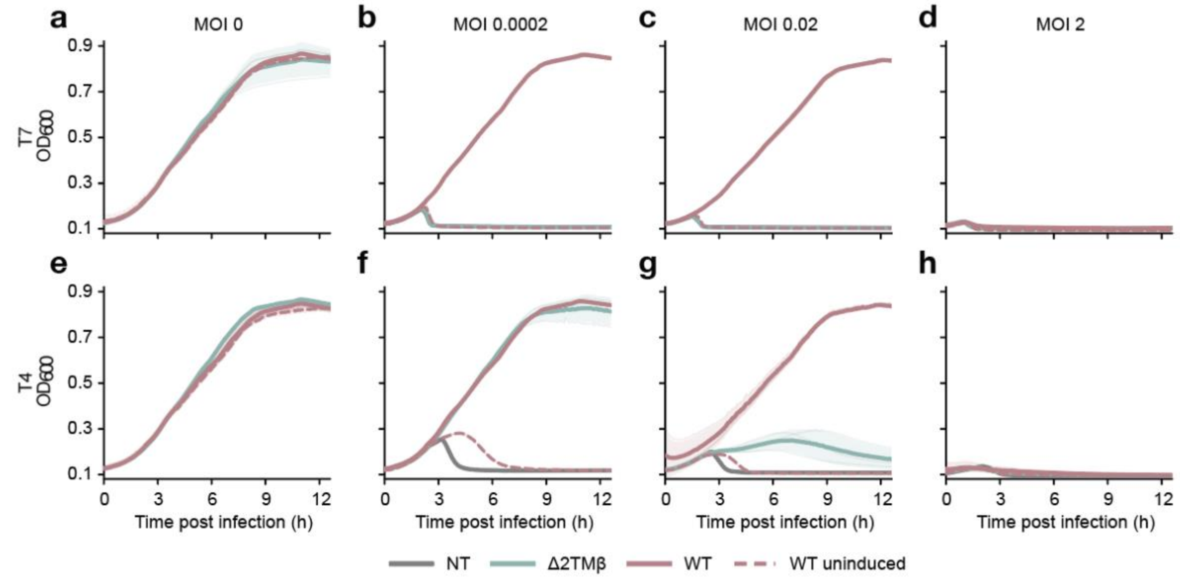


**Extended Data Fig. 3. *2TMβ-cas13b6* provides phage defense in liquid culture. a–d**, Growth of *E. coli* cultures expressing the indicated constructs following infection with phage T7 at multiplicities of infection (MOI) of 0 (a), 0.0002 (b), 0.02 (c) and 2 (d). **e–h**, Growth of *E. coli* cultures following infection with phage T4 at MOIs of 0 (e), 0.0002 (f), 0.02 (g) and 2 (h). Dashed lines indicate WT uninduced cultures. Data are plotted as $OD_{600}$ over time; shaded regions indicate s.d. across biological replicates.

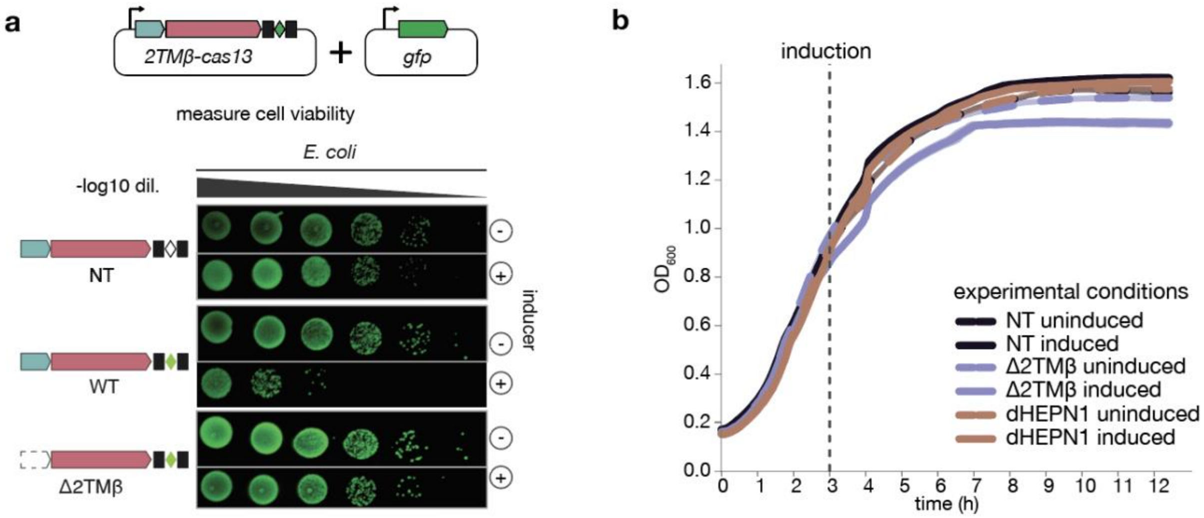


**Extended Data Fig. 4 | Target RNA expression triggers 2TMβ-dependent growth inhibition. a,** Schematic of the heterologous GFP-targeting assay and serial-dilution spot assays. *E. coli* cells were co-transformed with a plasmid encoding the *2TMβ–cas13b6* module programmed with a GFP-targeting spacer and a second plasmid expressing GFP. Target RNA expression was induced to activate Cas13 in the absence of phage infection. Cell viability was assessed for the non-targeting control (NT), WT targeting system and targeting system lacking 2TMβ (Δ2TMβ), without (−) or with (+) inducer. **b,** 12h post-seeding growth curves of phage-free GFP-targeting assay in *E. coli* carrying the *2TMβ–cas13* locus with a NT guide, the Δ*2TMβ–cas13* locus, or a *2TMβ–cas13*[dHEPN1] locus, alongside a *gfp* target plasmid. Solid and dotted lines denote induced and uninduced cultures, respectively. dHEPN1: Cas13 $R^{96}NFYSH^{101}$ > $A^{96}NFYSA^{101}$.

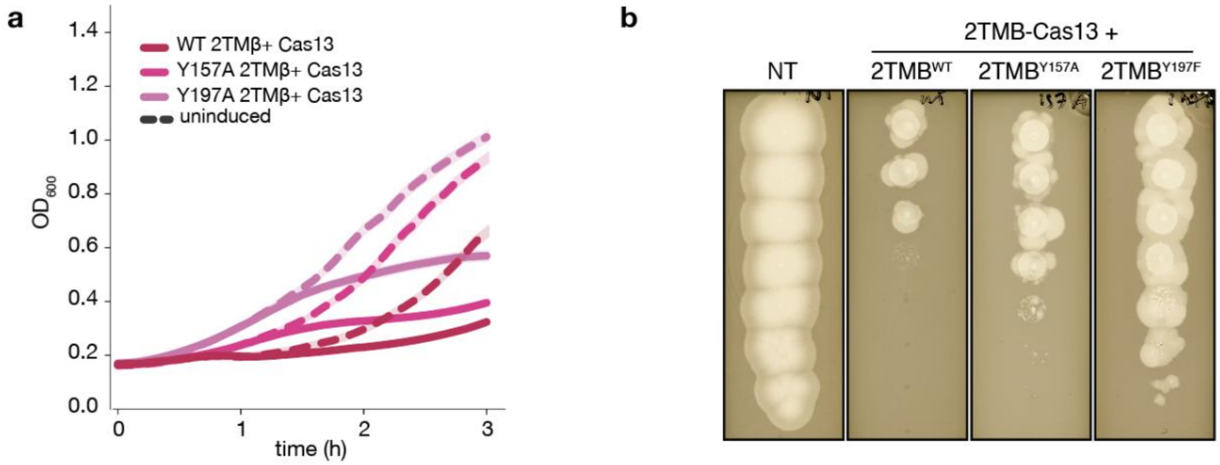


**Extended Data Fig. 5 | 2TMβ pocket mutants display a range of phenotypes upon system activation in response to target RNA expression from a plasmid or phage. a**, Growth curves of phage-free GFP-targeting assay in *E. coli* carrying the WT 2TMβ or 2TMβ pocket mutants Y157A and Y197A. Solid and dashed lines indicate induced and uninduced cultures, respectively. Growth was monitored by optical density at 600 nm ($OD_{600}$). **b**, Spot assays of phage T7 on *E. coli* co-expressing the WT 2TMβ-Cas13 operon and T7 targeting spacer, alongside either a WT or pocket mutant of 2TMβ. NT denotes the non-targeting control; targeting conditions contain WT 2TMβ or the Y157A or Y197F variant.

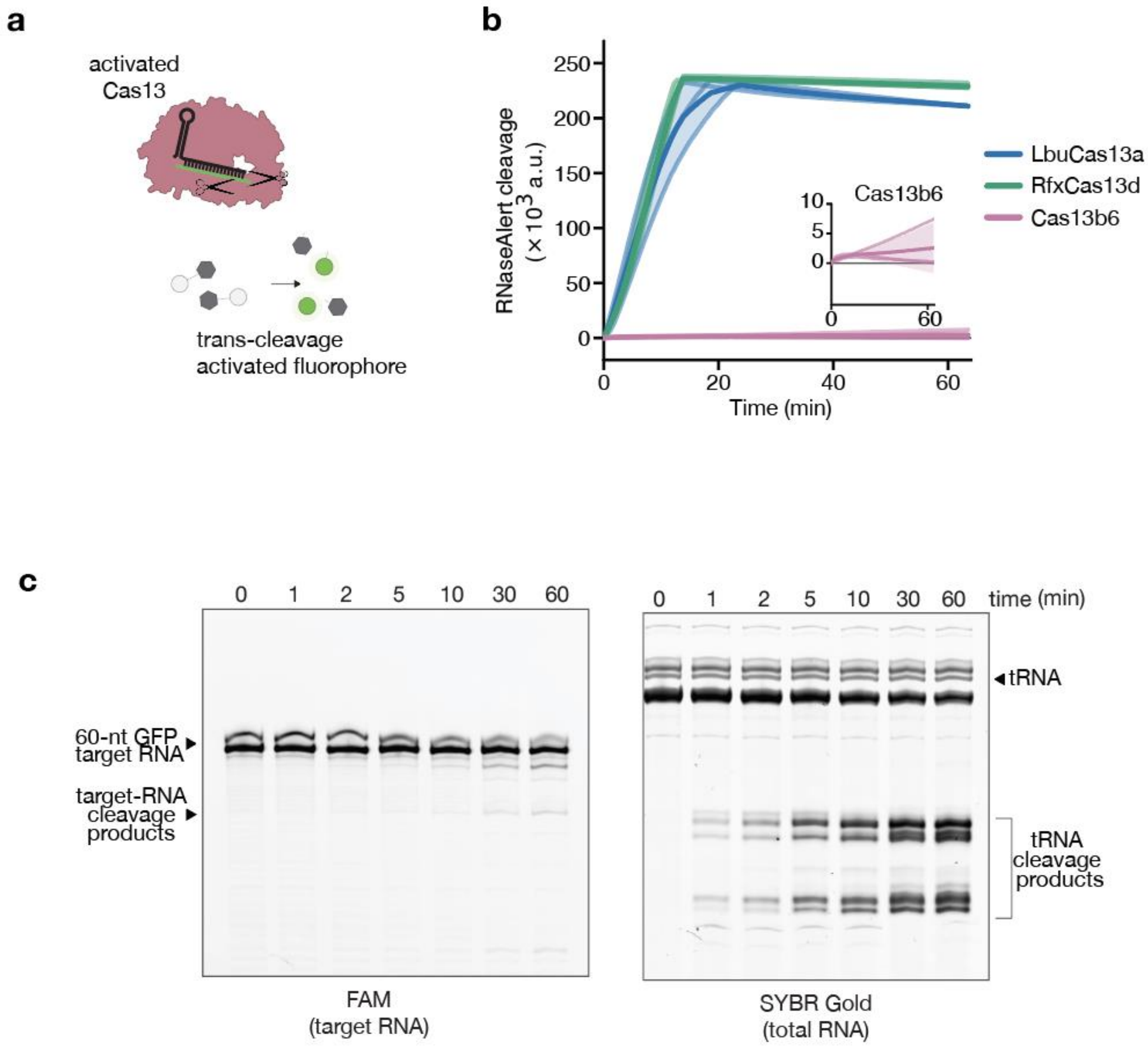


**Extended Data Fig. 6 | Cas13b6 shows minimal *cis-* and *trans-* cleavage on non-tRNA substrates.** **a,** Schematic of the collateral-cleavage assay used to compare trans-RNase activity of activated Cas13 enzymes. Target recognition by the Cas13–guide complex was monitored by cleavage of a fluorogenic FAM-5′-UUUU-3′ Iowa Black reporter. **b,** Time course of reporter cleavage by LbuCas13a, RfxCas13d and Cas13b6. Data are shown as mean ± s.d. from independent replicates. **c,** Denaturing gel analysis of target RNA incubated with the Cas13–guide RNA complexes. Positions of target RNA, and guide RNA are indicated.

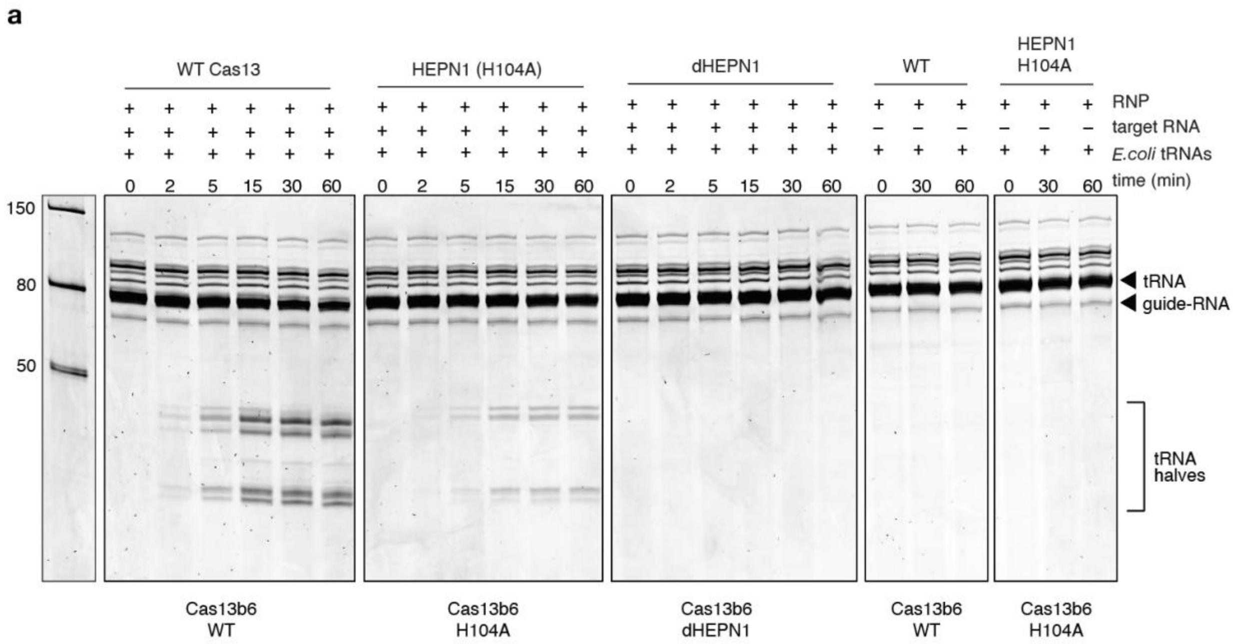


**Extended Data Fig. 7 | Cas13b6 cleavage of *E. coli* RNAs.** Denaturing gel analysis of *E. coli* RNA incubated with reconstituted Cas13b6 RNPs containing 500 nM Cas13 protein, 500 nM guide RNA and, where indicated, 100 nM target RNA. Reactions contained 1 µg/µL *E. coli* tRNAs and were sampled at the indicated times. Positions of intact tRNA, guide RNA and tRNA halves are indicated.

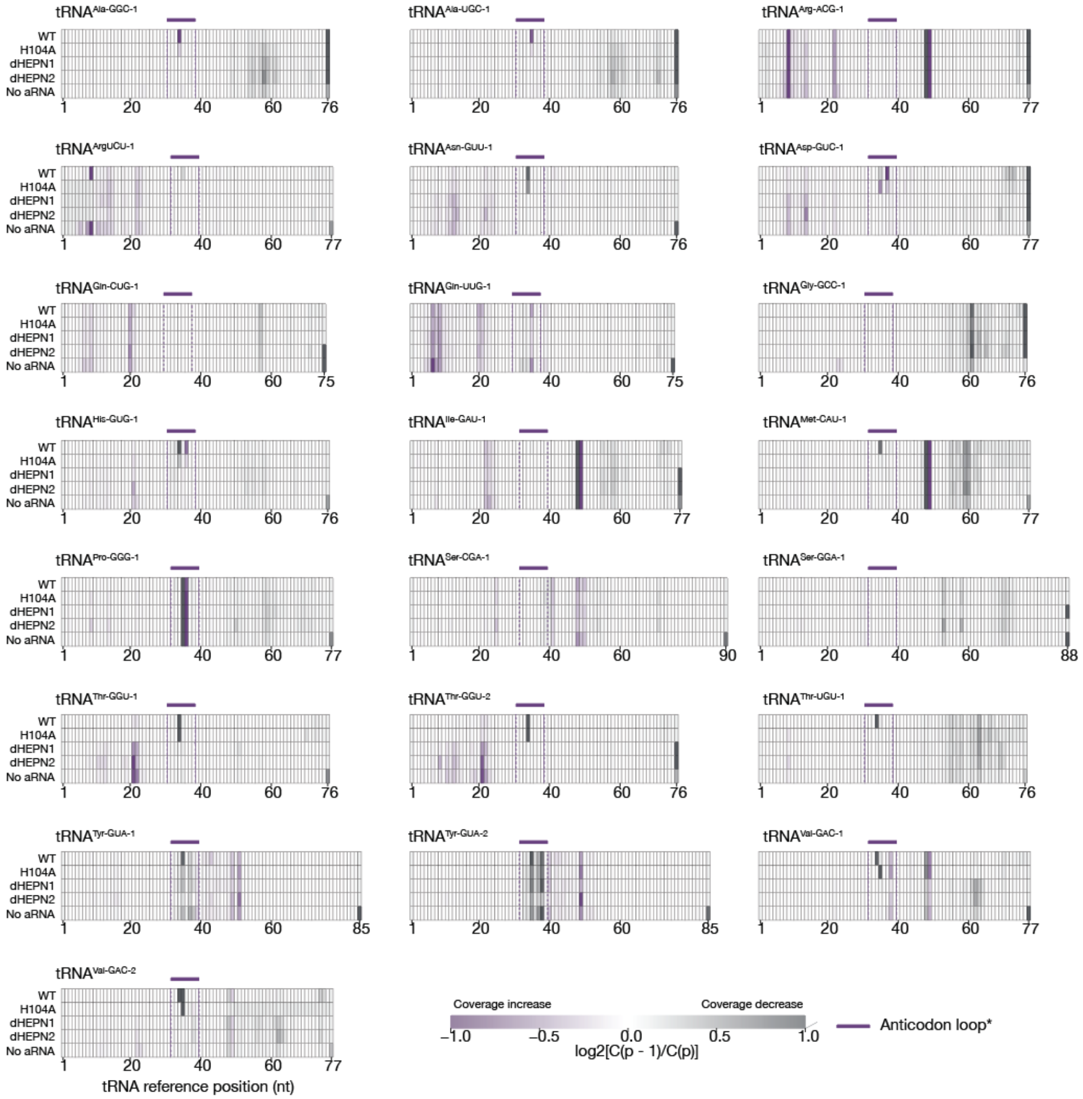


**Extended Data Fig. 8 | Small-RNA-seq profiles of read-coverage changes across tRNA reference sequences following Cas13b6 activation *in vitro.* a,** Heatmaps showing local changes in sequencing coverage across the complete reference sequences of 22 selected *E. coli* tRNAs. Rows represent reactions containing WT Cas13b6, H104A or the catalytically inactive mutants dHEPN1 and dHEPN2 in the presence of activator RNA, and the no-activator control (No aRNA). Values are arithmetic means of library-specific $\log_2[C(p-1)/C(p)]$ ratios, where C(p) is aligned-read coverage at reference position p. Two libraries were analyzed per condition, except dHEPN2, for which one library was available. Purple indicates increasing coverage and grey indicates decreasing coverage between adjacent positions; white indicates little or no change. Colours are capped at −1 and +1. Purple bars and dashed boundaries mark the anticodon region, operationally defined as anticodon-anchored positions 31–38 and mapped onto each reference sequence. The final x-axis tick indicates the reference length. Hatched cells denote undefined ratios or fewer than 20 reads at either adjacent position in any included library. dHEPN1: Cas13 $R^{96}NFYSH^{101} > A^{96}NFYSA^{101}$.

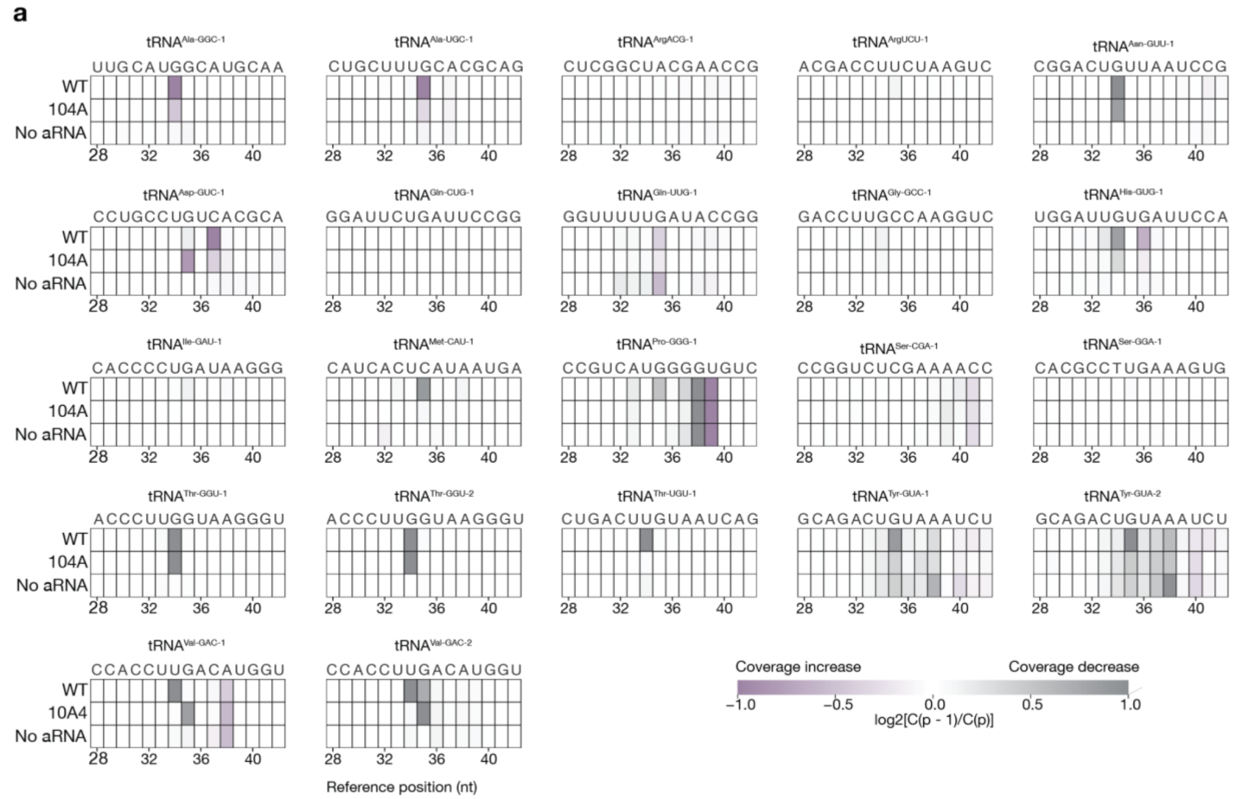


**Extended Data Fig. 9 | Local tRNA coverage changes following Cas13 activation *in vitro*. a,** Heatmaps showing adjacent-position changes in sequencing coverage across reference positions 28–42 of 22 selected *E. coli* tRNAs. Rows show WT Cas13b6 and H104A reactions with activator RNA, and the no-activator control (No aRNA). Values represent the arithmetic mean of library-specific $\log_2[C(p-1)/C(p)]$ ratios from two libraries per condition, where C(p) is aligned-read coverage at reference position p. Purple indicates increasing coverage and grey indicates decreasing coverage between adjacent positions; white indicates little or no change. Colours are capped at −1 and +1. Undefined ratios are shown in dark grey. Nucleotide sequences are displayed above each heatmap. Coordinates refer to the corresponding tRNA reference sequences, rather than standardized structural numbering.

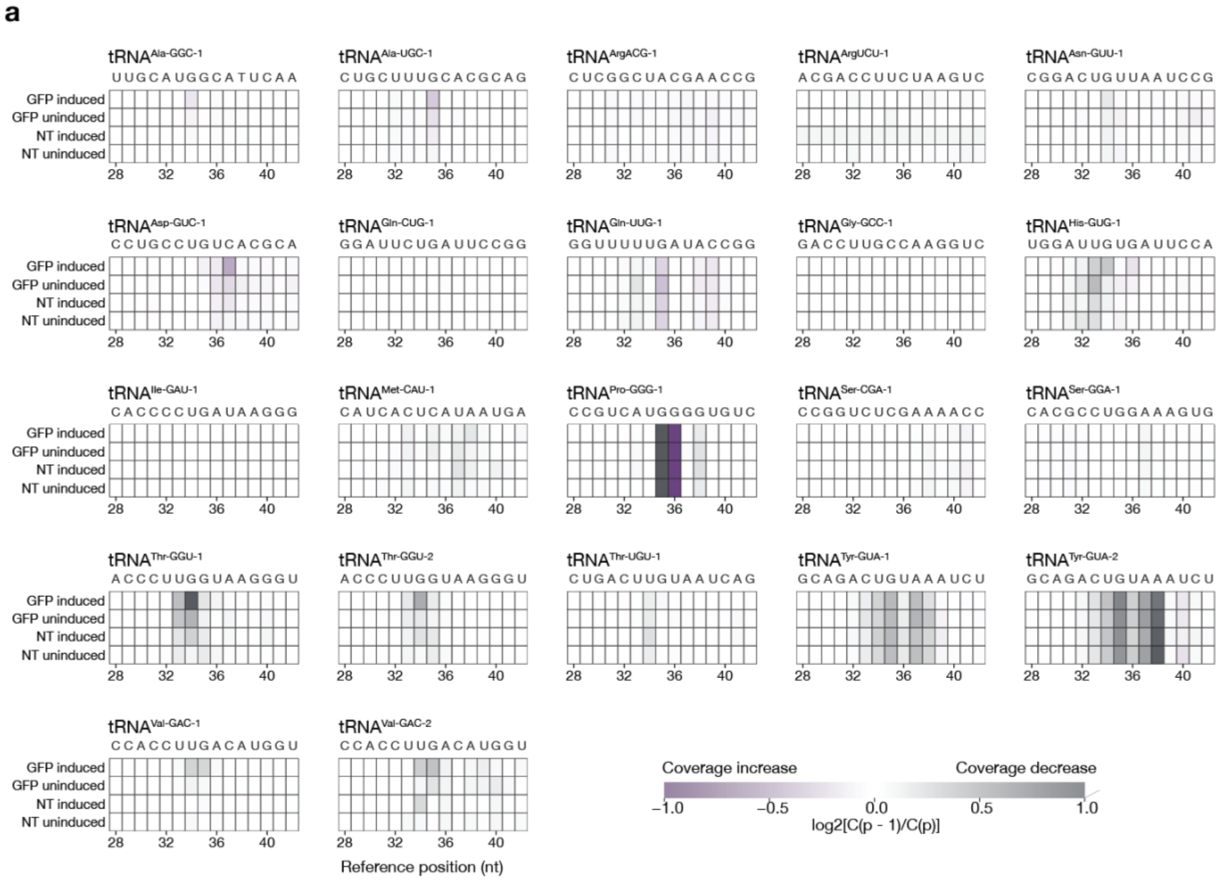


**Extended Data Fig. 10 | Small-RNA-seq profiles of tRNA anticodon-region coverage changes *in vivo*.** **a,** Heatmaps showing adjacent-position changes in sequencing coverage across reference positions 28–42 of 22 selected *E. coli* tRNAs. Rows represent cells carrying the 2TMβ–Cas13b6 system programmed with a GFP-targeting guide or a non-targeting guide (NT), under induced or uninduced conditions. Values are arithmetic means of two library-specific $\log_2[C(p-1)/C(p)]$ ratios per condition, where C(p) is aligned-read coverage at reference position p. Purple indicates increasing coverage and grey indicates decreasing coverage between adjacent positions; white indicates little or no change. Colours are capped at −1 and +1. Reference nucleotide sequences are displayed above each heatmap, and coordinates refer to the corresponding tRNA reference sequence rather than standardized structural numbering. Identical tRNA gene copies were pooled. Reads ending at the annotated mature tRNA 3′ terminus (A76) were excluded before coverage calculation. Ratios were displayed only when both adjacent positions had at least 20 reads in each library.

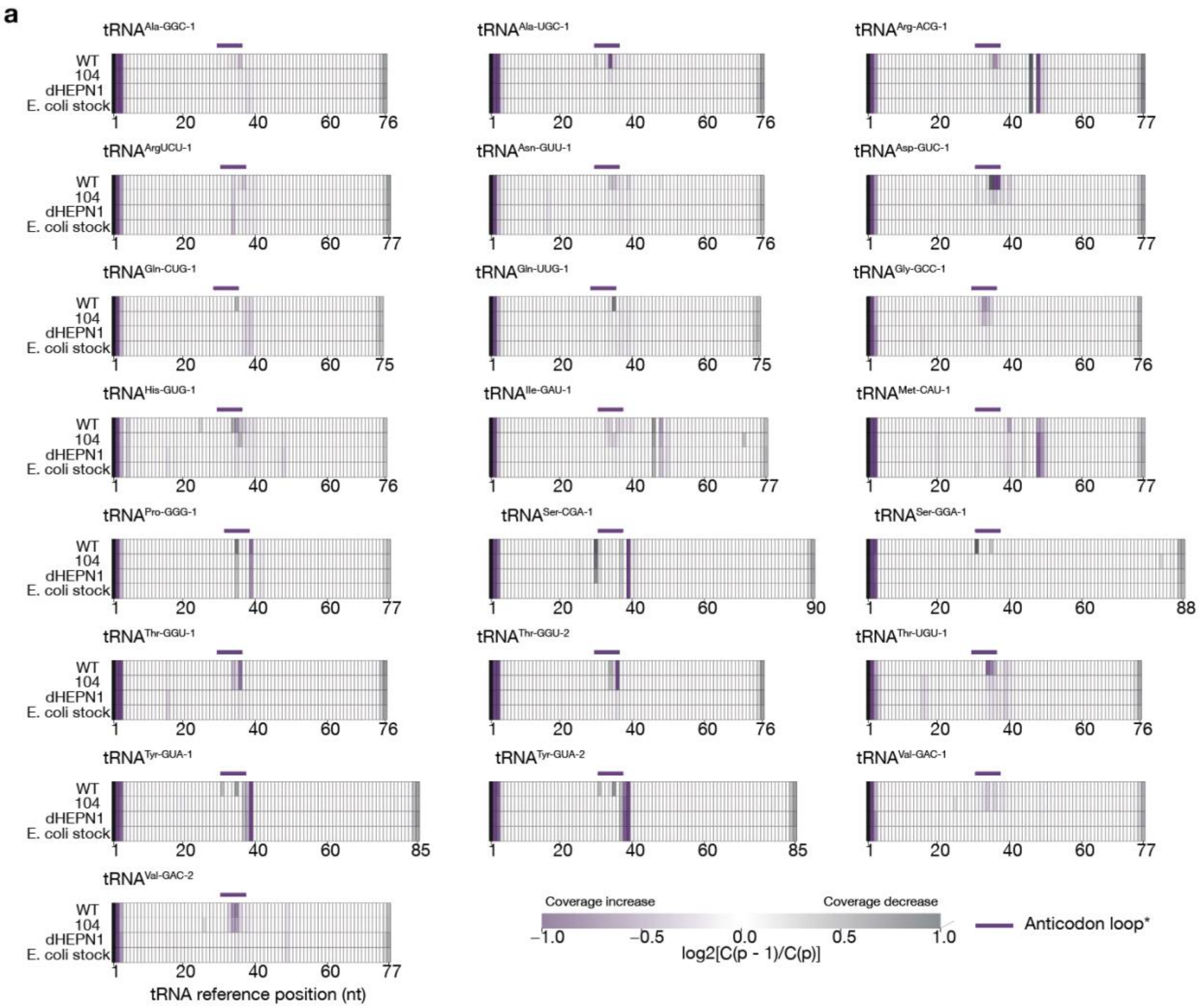


**Extended Data Fig. 11 | MSR-seq profiles of read-coverage changes across tRNA reference sequences. a,** Heatmaps showing local changes in sequencing coverage across the complete reference sequences of 22 selected *E. coli* tRNAs. Rows represent libraries from *in vitro* reactions containing WT Cas13b6, H104A or catalytically inactive dHEPN1, and the untreated *E. coli* tRNA stock. Each row represents one sequenced library. Values are calculated as $\log_2[C(p-1)/C(p)]$, where C(p) is read coverage at reference position p. Purple indicates increasing coverage and grey indicates decreasing coverage between adjacent positions; white indicates little or no change. Colours are capped at −1 and +1. Purple bars mark the anticodon region, operationally defined as anticodon-anchored positions 31–38 and mapped onto each reference sequence. The final x-axis tick indicates the reference length. Hatched cells denote undefined ratios or fewer than 20 reads at either adjacent position. dHEPN1: Cas13 $R^{96}NFYSH^{101}$ > $A^{96}NFYSA^{101}$

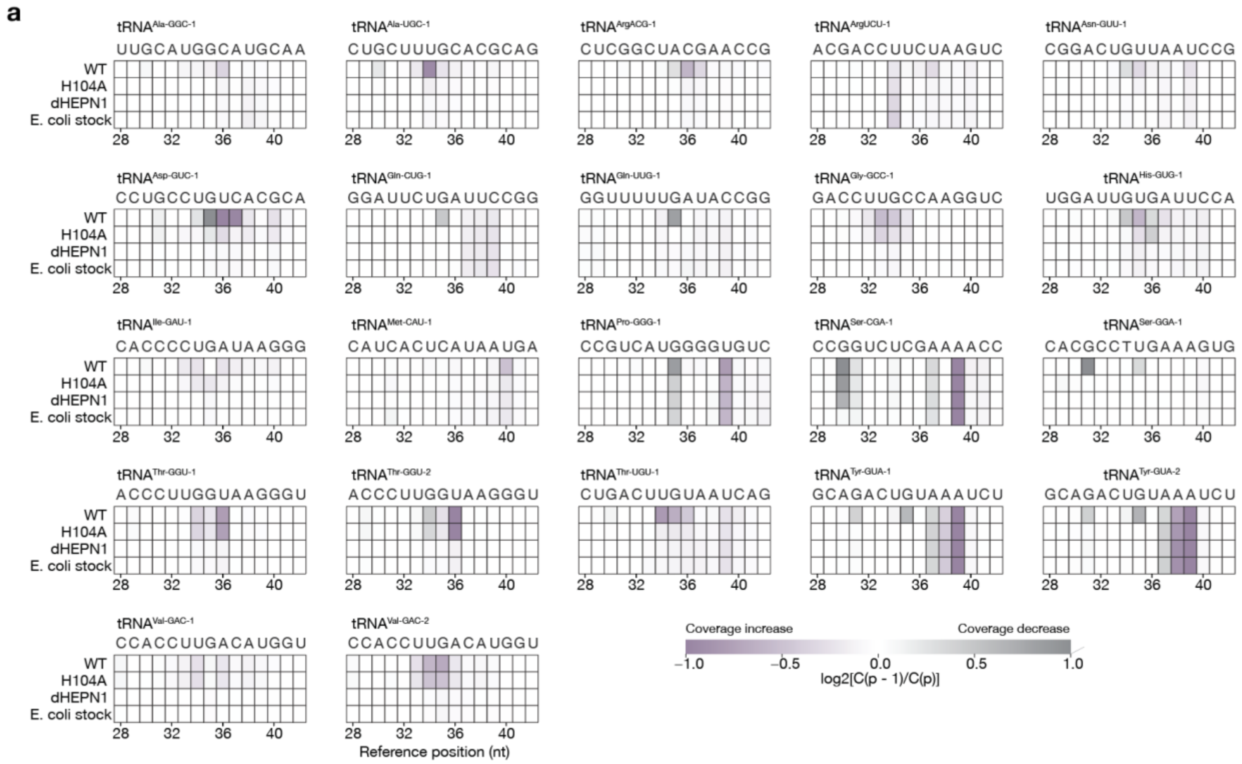


**Extended Data Fig. 12 | MSR-seq analysis of local coverage changes around tRNA anticodon regions. a,** Heatmaps showing adjacent-position changes in sequencing coverage across reference positions 28–42 of 22 selected *E. coli* tRNAs. Rows represent libraries from *in vitro* reactions containing WT Cas13b6, H104A or catalytically inactive dHEPN1, and the untreated *E. coli* tRNA stock. Each row represents one sequenced library. Values are calculated as $\log_2[C(p-1)/C(p)]$, where C(p) is read coverage at reference position p. Purple indicates increasing coverage and grey indicates decreasing coverage between adjacent positions; white indicates little or no change. Colours are capped at −1 and +1. Reference nucleotide sequences are displayed above each heatmap. Coordinates refer to the corresponding tRNA reference sequences rather than standardized structural numbering. dHEPN1: Cas13 $R^{96}NFYSH^{101}$ > $A^{96}NFYSA^{101}$.

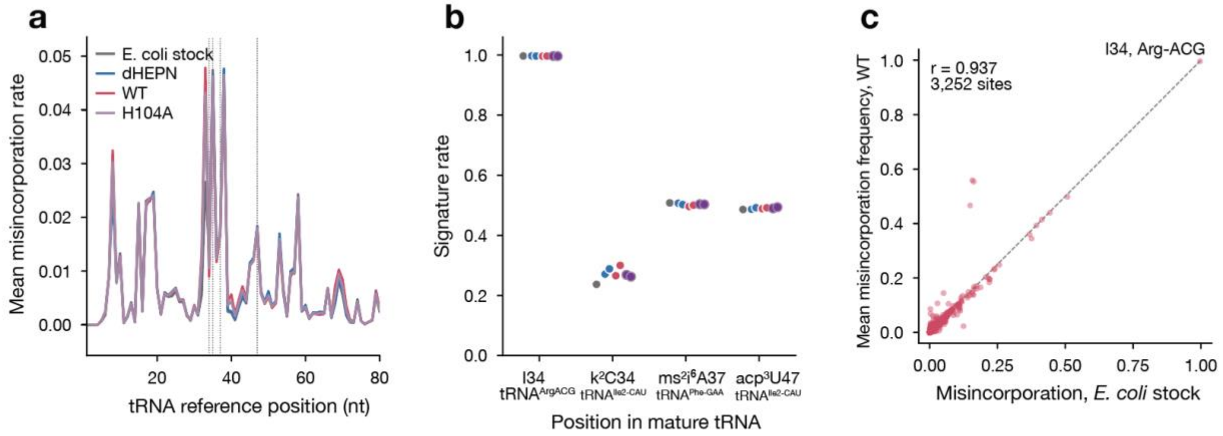


**Extended Data Fig. 13 | Modification-associated reverse-transcription signatures in MSR-seq libraries. a,** Mean misincorporation frequencies across tRNA reference positions in WT Cas13b6, H104A, catalytically inactive dHEPN1 and enzyme-free *E. coli* tRNA-stock libraries. Replicate libraries were averaged within each enzyme condition. Vertical dotted lines indicate reference positions corresponding to the diagnostic sites examined in **b**. **b,** Reverse-transcription signatures at sites associated with inosine (I34) in $tRNA^{Arg(AcG)}$, lysidine ($k^2C34$) and $acp^3U47$ in $tRNA^{Ile2(cAU)}$, and $ms^2i^6A37$ in $tRNA^{Phe(GAA)}$. Signature rates represent misincorporation frequencies for I34 and $ms^2i^6A37$, and deletion frequencies for $k^2C34$ and $acp^3U47$. Each point represents one library: two per enzyme condition and one tRNA-stock library. Colors correspond to **a**. Modification labels indicate expected assignments at the diagnostic sites. **c,** Comparison of mean WT misincorporation frequencies with the tRNA stock across 3,252 shared tRNA-position entries. Each point represents one entry; the dashed line indicates equality. Pearson r = 0.937. Positions in **a** are sequential reference coordinates; the Arg I34 site corresponds to reference position 35. These signatures support recovery of modified tRNAs but do not measure modification occupancy or exclude all reverse-transcription biases. dHEPN1: Cas13 $R^{96}NFYSH^{101}$ > $A^{96}NFYSA^{101}$.

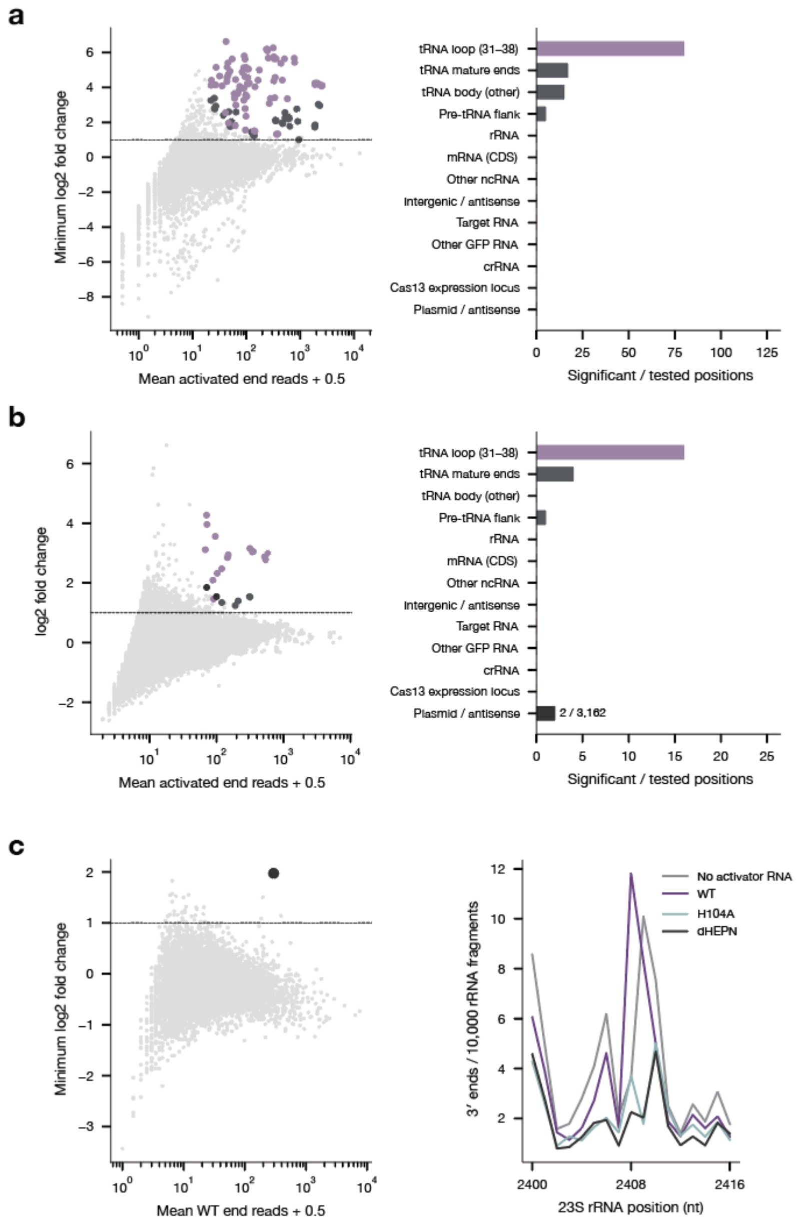


**Extended Data Fig. 14 | Activation-associated RNA termini are concentrated in tRNA anticodon regions. a,b,** Small-RNA-seq analysis of Cas13b6 reactions *in vitro* (**a**) and cellular RNA (**b**). Left, mean endpoint-read counts versus $\log_2$ fold change, taking the minimum across control comparisons where applicable. Purple marks significantly enriched tRNA anticodon-region endpoints; dark grey marks other significant endpoints. Dashed lines indicate $\log_2$ fold change = 1. Right, significant/tested positions by RNA class. **c,** Analysis of the rRNA-rich fraction (left) and normalized 23S rRNA 3′-end profiles surrounding the enriched endpoint U2408 (right). OFF, no activator; 104, H104A; dHEPN, pooled catalytically inactive controls. Enrichment required $\log_2$ fold change > 1 and Benjamini–Hochberg-adjusted $P < 0.05$ in negative-binomial models, relative to both no-activator and inactive controls (**a,c**) or non-targeting uninduced cells (**b**). Only positions with ≥20 summed reads were tested; 0/0 denotes no testable positions.

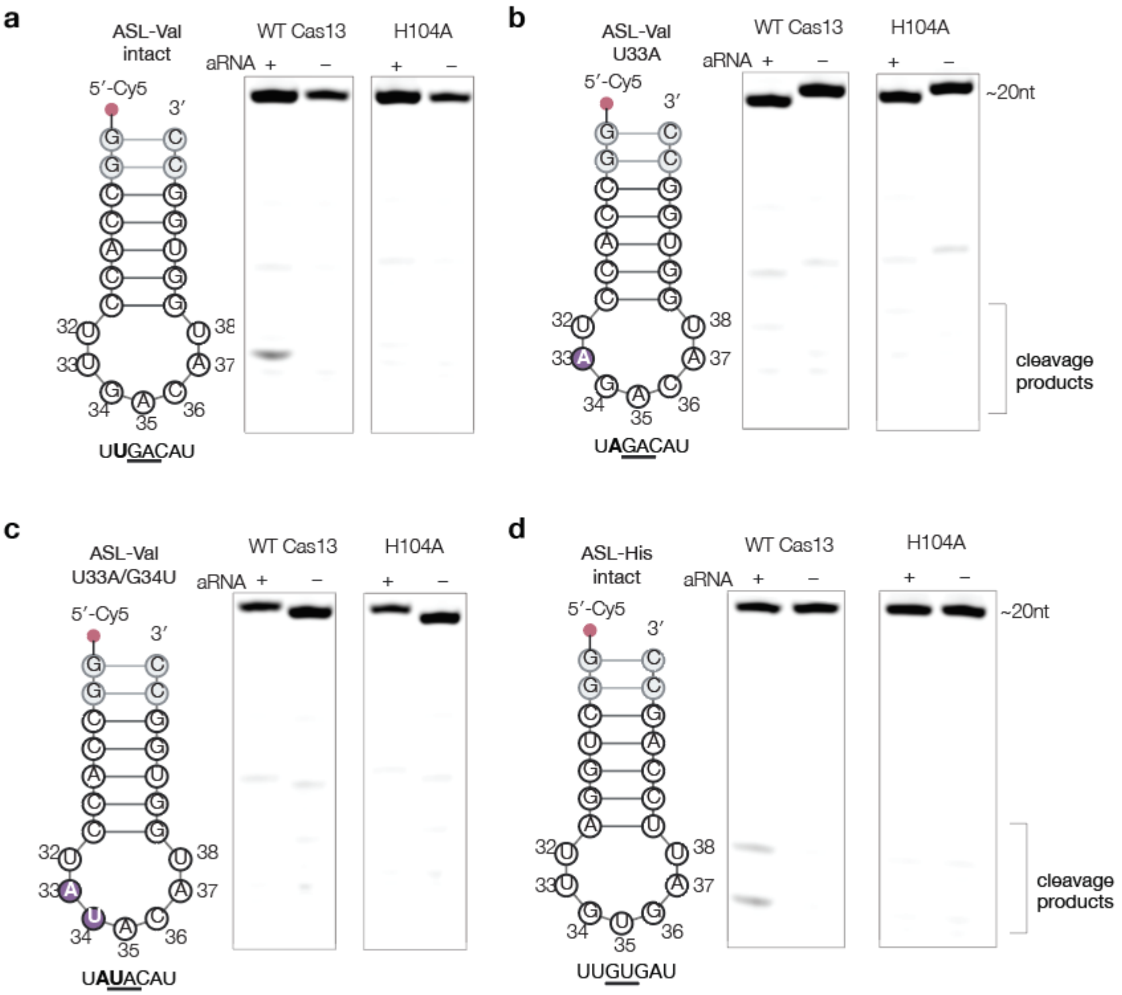


**Extended Data Fig. 15 | Cleavage of synthetic tRNA anticodon stem–loop substrates by Cas13b6.** **a–d,** Schematics and denaturing PAGE analysis of 5′-Cy5-labelled anticodon stem–loop (ASL) substrates corresponding to unmutated Val (**a**), Val U33A (**b**), Val U33A/G34U (**c**) and unmutated His (**d**). Substrates were incubated with WT Cas13b6 or H104A in the presence (+) or absence (−) of activator RNA (aRNA). Nucleotide positions follow conventional tRNA numbering; purple nucleotides indicate substitutions, and underlining identifies the anticodon triplet at positions 34-36. Intact substrates and products retaining the 5′-Cy5 label were detected by fluorescence imaging. Experiments were performed in duplicate.

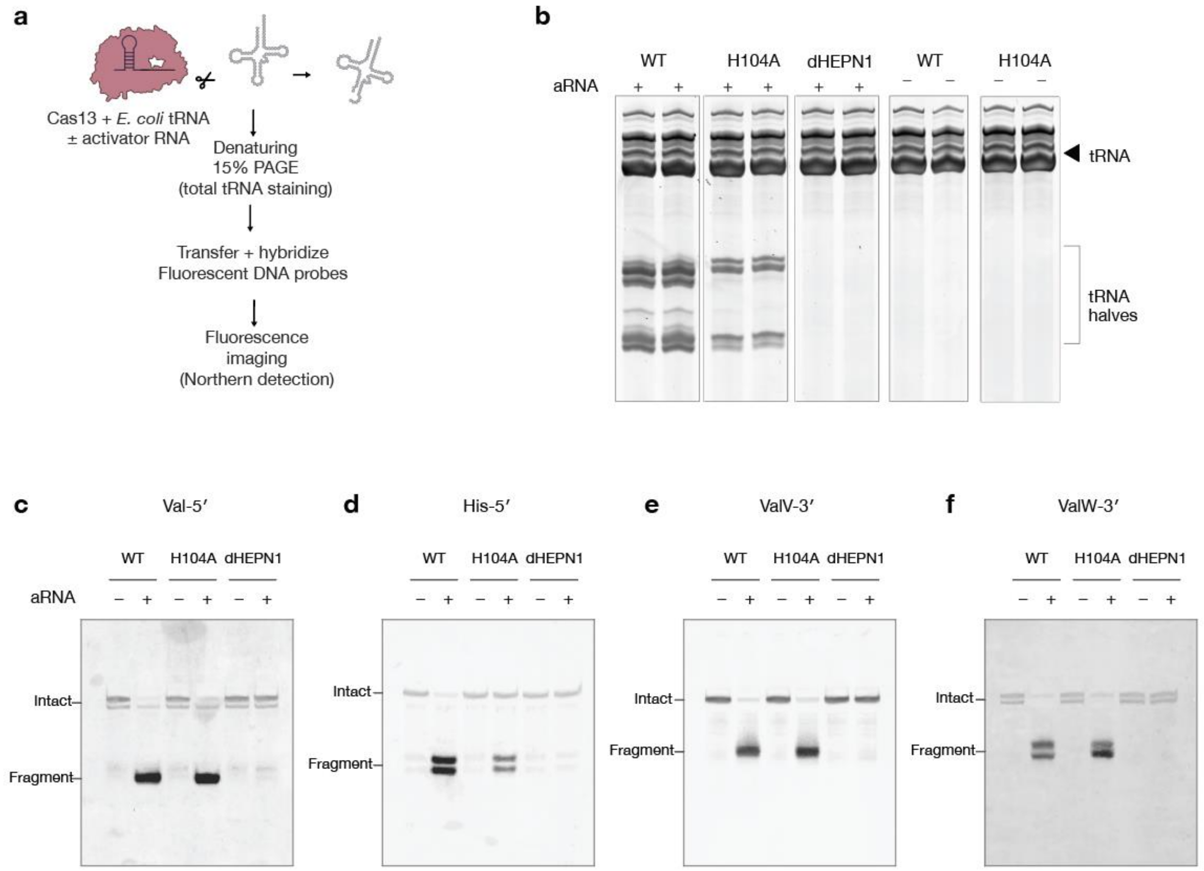


**Extended Data Fig. 16 | Activator-dependent cleavage of tRNAs by Cas13. a,** Experimental workflow. Purified *E. coli* tRNA was incubated with Cas13 variants with or without activator RNA (aRNA), followed by denaturing 15% PAGE and total-RNA staining or Northern blotting. **b,** Total-RNA staining of reactions containing WT Cas13, H104A or catalytically inactive dHEPN1, with (+) or without (−) aRNA. Paired lanes show replicate samples; lane groups are displayed separately. **c–f,** Northern blots analysis of the same RNA samples with Cy5-labelled DNA oligonucleotides complementary to the 5′ regions shared by $tRNA^{Val}_{GAC}$ ValV and ValW(Val-5′; **c**), the 5′ region $tRNA^{His}_{GUG}$ (His-5′; **d**), and the 3′ regions of the tRNAs ValV (ValV-3′; **e**), and ValW (ValW-3′; **f**). Intact-sized tRNAs and cleavage fragments are indicated. dHEPN1: Cas13 $R^{96}NFYSH^{101} > A^{96}NFYSA^{101}$.

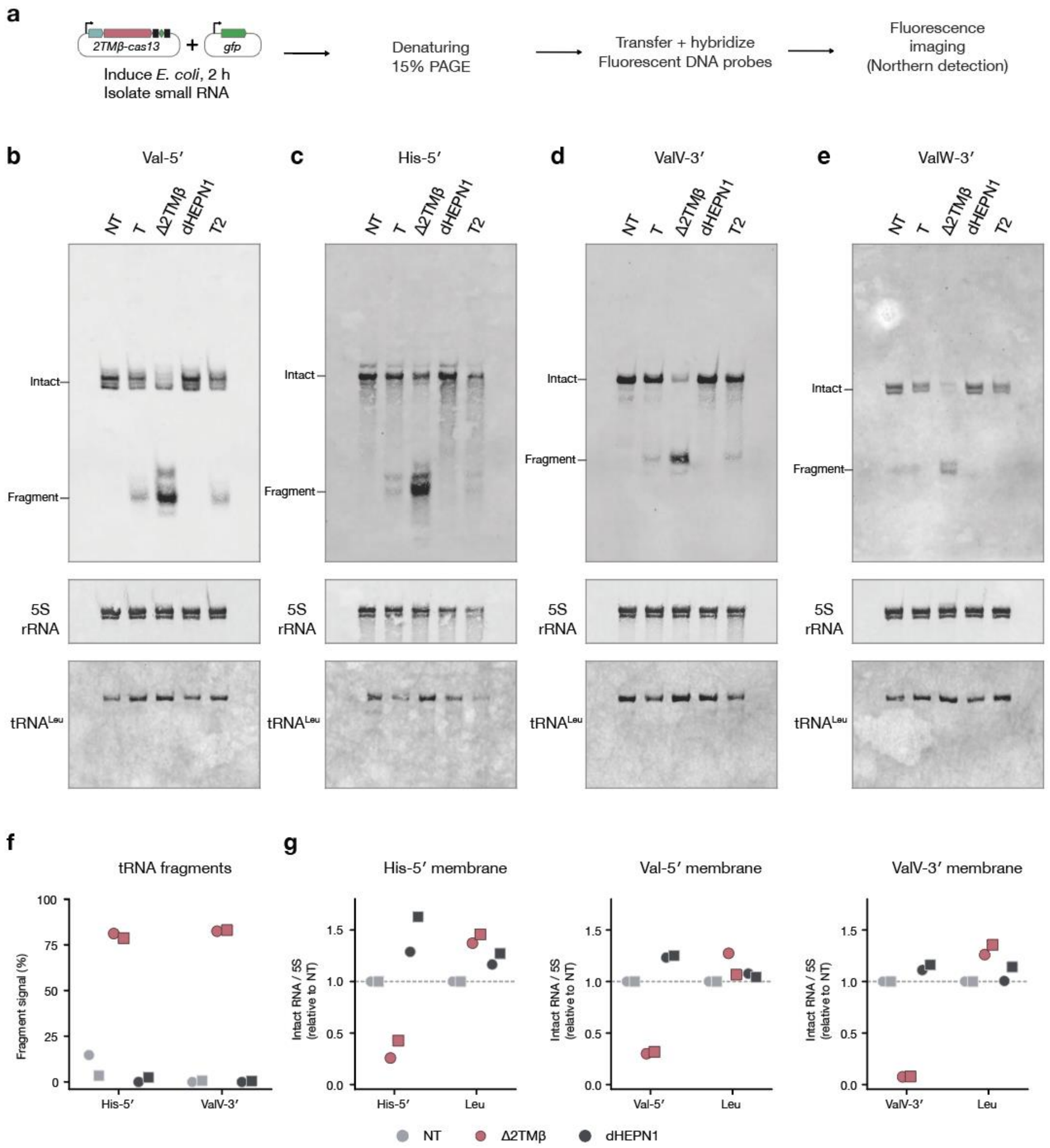


**Extended Data Fig. 17 | Northern blot detection of Cas13-dependent tRNA cleavage *in vivo*. a,** Experimental workflow. RNA was isolated from *E. coli* 2 h after induction, resolved by denaturing 15% PAGE and analyzed by Northern blotting. **b–e,** Northern blots detected using Cy5-labelled DNA probes complementary to the shared 5′ region of ValV and ValW tRNAs (Val-5′; **b**), the 5′ region of His tRNA (His-5′; **c**) and the respective 3′ regions of ValV (ValV-3′; **d**) and ValW (ValW-3′; **e**). NT denotes the non-targeting control; T1 and T2 denote two independent biological replicates of the WT targeting condition. Δ2TMβ and catalytically inactive dHEPN1 were examined under targeting conditions. Intact-sized tRNAs and cleavage fragments are indicated. Corresponding loading and specificity controls are shown with Cy3-labelled DNA oligonucleotides complementary to the 5S rRNA and tRNA$^{Leu}$ beneath each panel. **f,** Semi-quantitative analysis of fragment-associated signal detected with the His-5′ and ValV-3′ probes. Percentages were calculated as 100 × fragment signal/(intact + fragment signal), using background-corrected intensities. **g,** Intact substrate-tRNA and tRNA$^{Leu}$ control signals normalized to the corresponding 5S rRNA signal and expressed relative to the non-targeting control within each membrane and replicate set. Each plot shows the indicated substrate probe and the $tRNA^{Leu}$ control measured on the same membrane. Light grey, NT; pink, Δ2TMβ; dark grey, dHEPN1. Circles and squares indicate replicate sets 1 and 2, respectively; dashed lines indicate NT = 1. WT targeting lanes were excluded from quantification. Val-5′ fragment signals were excluded from **f** because of saturation, whereas its unsaturated intact signal was retained in **g**. ValW-3′ measurements were excluded from **f–g** owing to uneven background. $tRNA^{Leu}$ measurements across membranes are control reprobes, not additional biological replicates. These measurements represent detected signal fractions and relative intact-RNA abundance, rather than calibrated percentages of cellular tRNA molecules cleaved. dHEPN1: Cas13 $R^{96}NFYSH^{101}$ > $A^{96}NFYSA^{101}$.

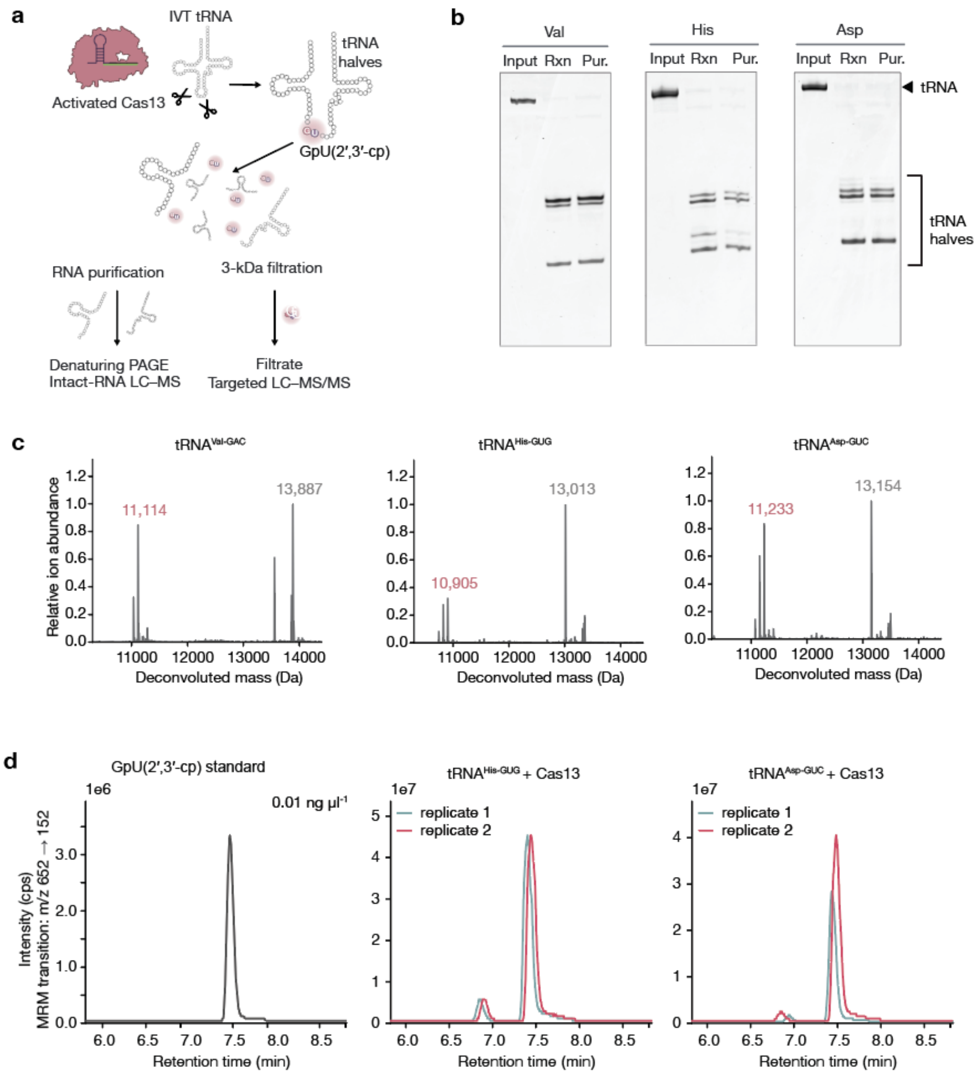


**Extended Data Fig. 18 | Characterization of Cas13 cleavage products from *in vitro*-transcribed tRNAs. a,** Experimental workflow. In vitro-transcribed (IVT) Val, His and Asp tRNAs were incubated with activated Cas13. RNA products were purified for denaturing PAGE and intact-RNA liquid chromatography–mass spectrometry (LC–MS). Parallel preparations were passed through a 3-kDa molecular-weight-cut-off filter, and the filtrates were analyzed by targeted LC–MS/MS. GpU(2′,3′-cp) formation is illustrated for the His and Asp substrates. **b,** Denaturing PAGE of IVT tRNA substrates before cleavage (Input), Cas13 cleavage reactions before purification (Rxn) and purified RNA submitted for intact-RNA LC–MS (Pur.). Substrate tRNA and cleavage-fragment regions are indicated. **c,** Deconvoluted mass spectra of purified products from Val, His and Asp tRNA cleavage reactions. Labelled masses, in daltons, correspond to assigned 5′ fragments (pink) and 3′ fragments (grey). Each spectrum is normalized independently; relative ion abundance does not represent the fraction of substrate cleaved. **d,** Targeted LC–MS/MS chromatograms of an authentic GpU(2′,3′-cp) standard (0.01 ng $\mu l^{-1}$) and filtrates from His and Asp cleavage reactions, monitored using the multiple-reaction-monitoring transition *m/z* 652 → 152. Coloured traces show two technical replicates per substrate. GpU(2′,3′-cp) denotes a GU dinucleotide bearing a terminal uridine 2′,3′-cyclic phosphate.

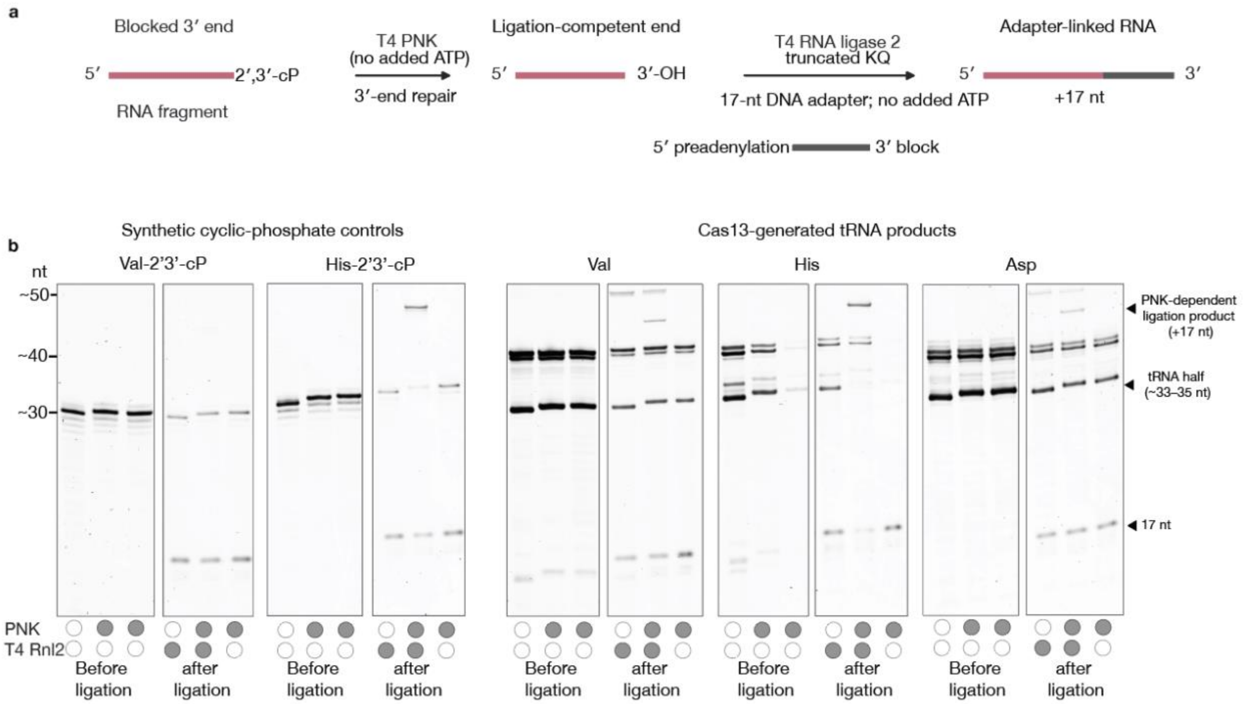


**Extended Data Fig. 19 | PNK-dependent adapter ligation of Cas13-generated tRNA fragments. a,** Experimental workflow. RNA fragments bearing blocked 3′ ends were treated with T4 polynucleotide kinase (PNK) without added ATP to generate ligation-competent 3′-OH ends. A 5′-preadenylated, 3′-blocked, 17-nt DNA adapter was subsequently ligated using T4 RNA ligase 2 truncated KQ without added ATP. Adapter attachment increases product length by 17 nucleotides. **b,** Denaturing PAGE analysis of synthetic Val and His RNA controls bearing terminal 2′,3′-cyclic phosphates and Cas13-generated Val, His and Asp tRNA products. For each substrate, the left and right lane groups show samples before and after the ligation step, respectively. Filled and open circles indicate inclusion and omission of the indicated enzymes. The ligation-stage samples include a no-PNK control and a no-ligase control. Positions of unligated tRNA halves, adapter-linked products and the free 17-nt adapter are indicated.

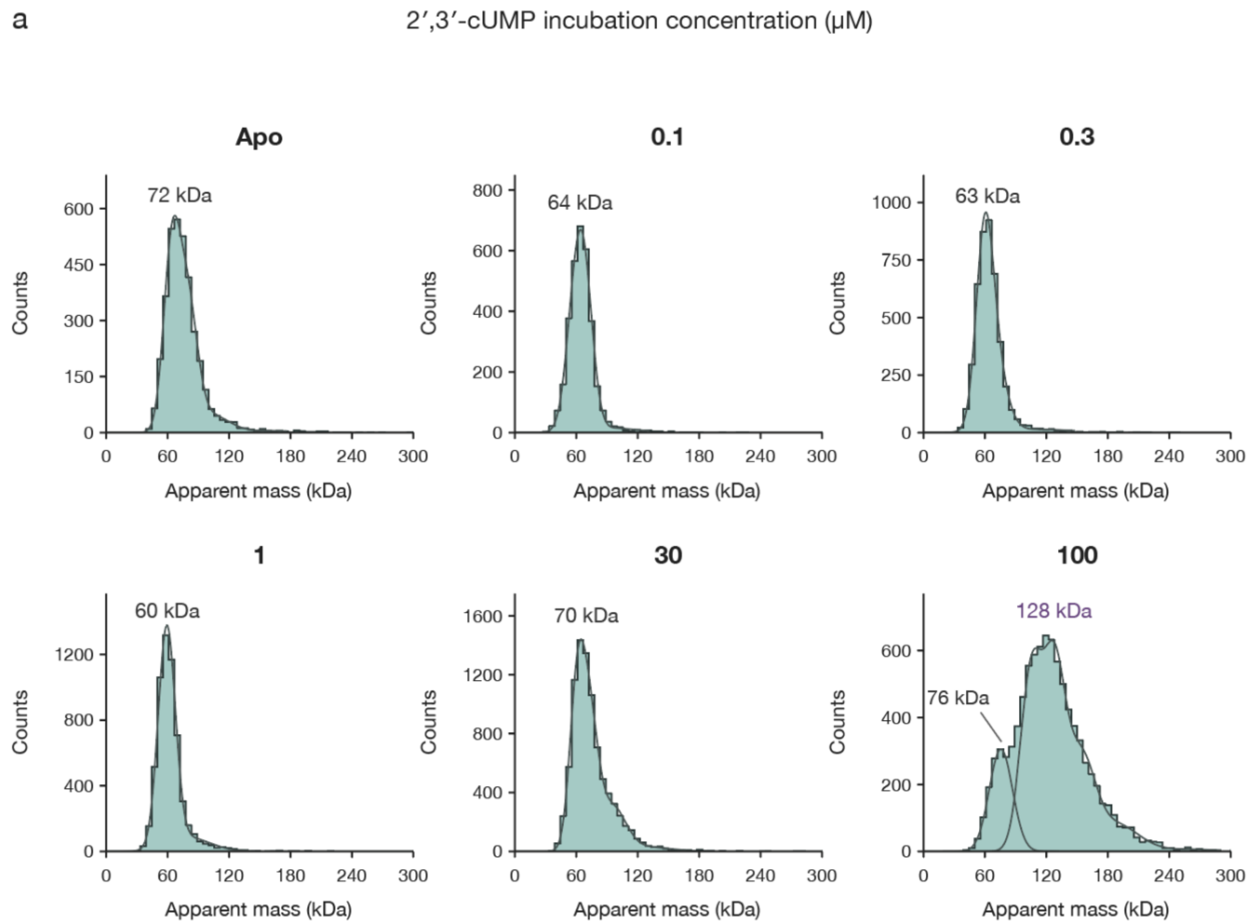


**Extended Data Fig. 20 | Mass-photometry analysis of the 2TMβ β-barrel across a 2′,3′-cUMP titration. a,** Apparent mass distributions of purified WT MBP–2TMβ β-barrel (1 μM) incubated without ligand (apo) or with the indicated concentrations of 2′,3′-cUMP. Concentrations shown refer to the incubation mixtures. Samples were diluted 20-fold into ligand-free buffer before measurement, yielding 50 nM protein and proportionally diluted ligand concentrations. Histograms show detected particle counts, with fitted distributions overlaid; annotations indicate apparent molecular masses. Following incubation with 100 μM 2′,3′-cUMP (5 μM during measurement), a higher-mass population near 128 kDa appeared alongside a population near 76 kDa. At least two replicate measurements were acquired per sample.

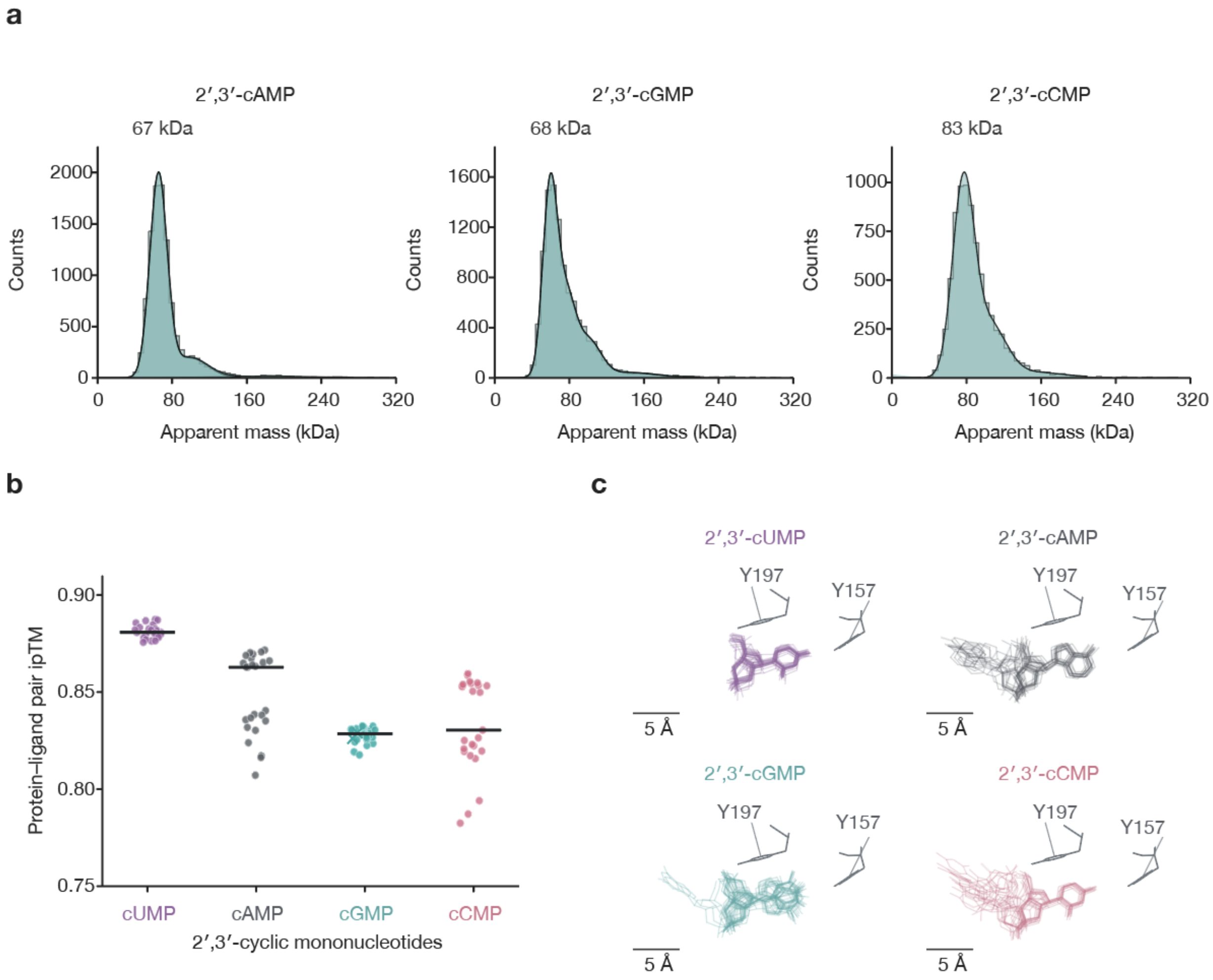


**Extended Data Fig. 21 | Predicted cyclic-nucleotide recognition and mass-photometry analysis of 2TMβ. a,** Mass-photometry distributions of purified soluble WT MBP–2TMβ β-barrel incubated with 2′,3′-cCMP, 2′,3′-cAMP or 2′,3′-cGMP. Labels indicate the apparent masses of the dominant populations. **b,** Protein–ligand pair ipTM scores from Protenix predictions containing 2TMβ and the indicated nucleoside 2′,3′-cyclic monophosphates. U>p, A>p, G>p and C>p denote 2′,3′-cUMP, 2′,3′-cAMP, 2′,3′-cGMP and 2′,3′-cCMP, respectively. Each point represents the mean score of the two protein–ligand pairs within one prediction. Twenty-five predictions were generated per ligand using five seeds and five samples per seed; horizontal bars indicate medians. **c,** Superposition of the 50 ligand instances per condition after alignment of protein Cα atoms at residues 81–214. Ligands are coloured as in **b**; Y157 and Y197 from the reference cUMP model are shown in grey. All four views use the same orientation and scale. Scale bars, 5 Å. Model confidence and pose consistency are descriptive computational measures, not measurements of binding affinity.

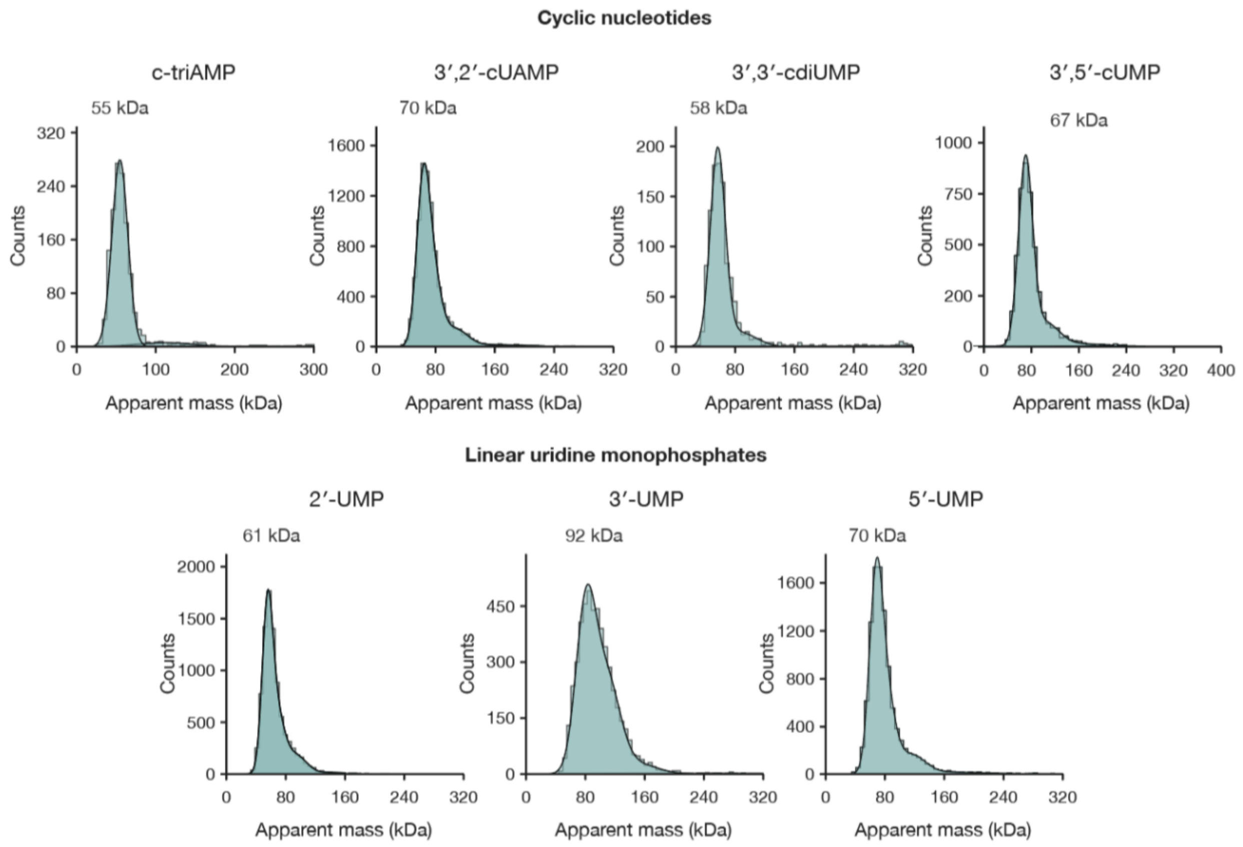


**Extended Data Fig. 22 | Mass-photometry analysis of the 2TMβ β-barrel with related cyclic and linear nucleotides. a,** Apparent mass distributions of purified WT MBP–2TMβ β-barrel incubated with cyclic tri-AMP (c-triAMP), 3′,2′-cyclic UMP–AMP (3′,2′-cUAMP), 3′,3′-cyclic di-UMP (3′,3′-cdiUMP), 3′,5′-cUMP, or uridine 2′-, 3′- or 5′-monophosphate. Incubation mixtures contained 1 μM protein and 300 μM of the indicated ligand. Samples were diluted 20-fold into ligand-free buffer before measurement, yielding 50 nM protein and 15 μM ligand. Histograms show detected particle counts; annotations indicate apparent masses of the dominant populations. At least two replicate measurements were acquired per sample.

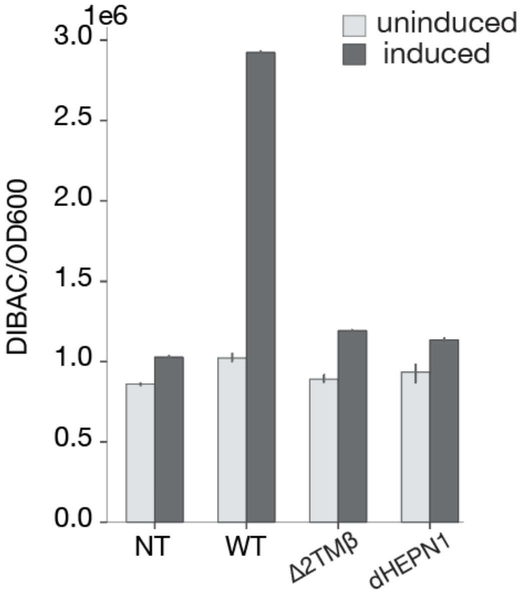


**Extended Data Fig. 23 | DiBAC fluorescence following activation of the 2TMβ–Cas13b6 system. a,** DiBAC fluorescence normalized to $OD_{600}$ in *E. coli* carrying the complete system (WT), a 2TMβ deletion (Δ2TMβ) or the Cas13b6 HEPN1 catalytic mutant (dHEPN1), under induced and uninduced GFP-targeting conditions. NT, non-targeting guide control. dHEPN1: Cas13 $R^{96}$NFYSH$^{101}$ > $A^{96}$NFYSA$^{101}$.

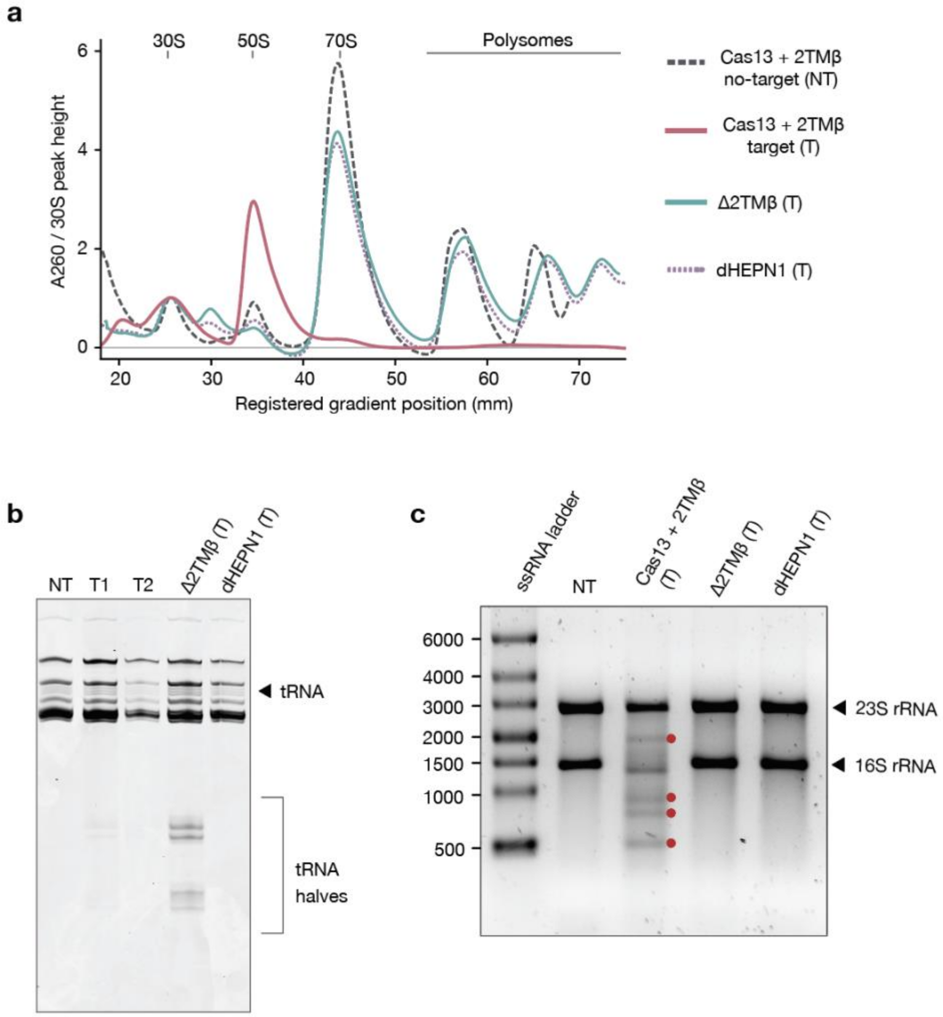


**Extended Data Fig. 24 |** Ribosome profiles and rRNA integrity following Cas13-2TMβ activation. **a**, Sucrose-gradient polysome profiles of *E. coli* harvested 2 h after induction under non-targeting (NT) or targeting (T) conditions. Absorbance at 260 nm was normalized to the 30S peak height, and gradient positions were registered across samples. Assigned 30S, 50S and 70S peaks and the polysome region are indicated. Δ2TMβ and dHEPN1 were analyzed under targeting conditions. **b**, Gel analysis of cellular RNA showing tRNAs and smaller products in the tRNA-half size range. T1 and T2 denote targeting samples. **c**, Gel analysis of total cellular RNA showing the indicated 23S and 16S rRNA bands; the single stranded RNA ladder is labelled. Red dots indicate additional RNA species present following Cas13 activation, likely corresponding to the degradation of full length rRNA species. Δ2TMβ, deletion of the 2TMβ effector; dHEPN1, catalytically inactive Cas13 HEPN1 mutant (Cas13 $R^{96}NFYSH^{101}$ > $A^{96}NFYSA^{101}$).

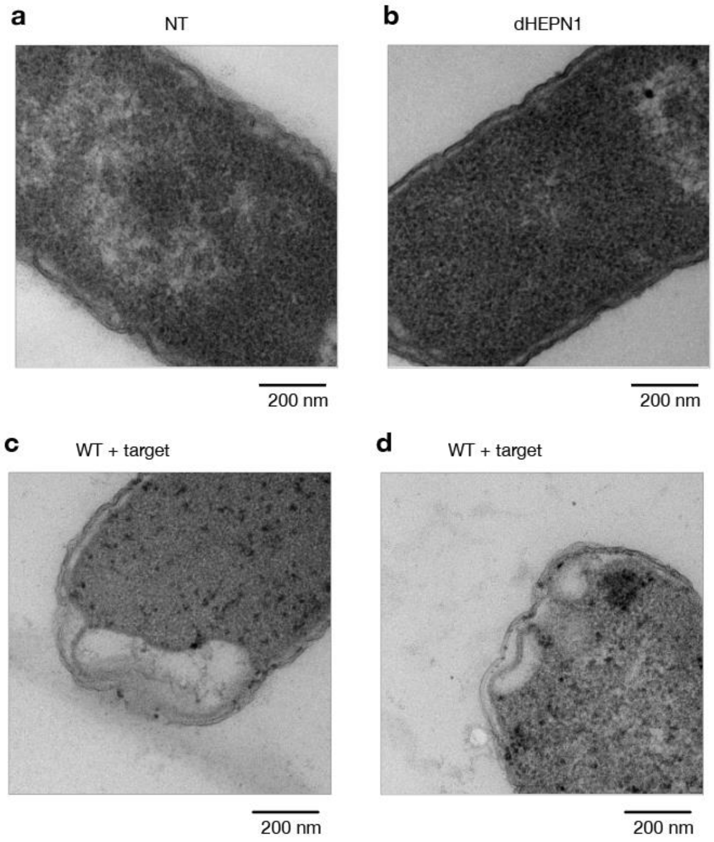


**Extended Data Fig. 25 | Ultrastructural changes associated with activation of the 2TMβ–Cas13 system. a,b,** Transmission electron microscopy images of *Escherichia coli* carrying the non-targeting control (NT; **a**) or the catalytically inactive dHEPN1 system (**b**). **c,d,** Selected images of cells carrying the WT system under targeting conditions, showing polar electron-lucent spaces and irregular cell-envelope contours. Images derive from one culture per condition; **c** and **d** show different fields from the same WT grid. These images provide qualitative examples of ultrastructural changes rather than measurements of phenotype frequency. Scale bars, 200 nm. dHEPN1: Cas13 $R^{96}NFYSH^{101}$ > $A^{96}NFYSA^{101}$.
